# Biogeography and cryptic diversity of the ancient centipede genus *Digitipes* (Scolopendromorpha) in South and Southeast Asia

**DOI:** 10.64898/2026.07.28.741128

**Authors:** Payal Dash, Pragyadeep Roy, Jahnavi Joshi

## Abstract

Understanding the relative roles of vicariance and dispersal in shaping diversity and distribution patterns is central to historical biogeography. In this study, we investigate the historical biogeography of the ancient centipede genus *Digitipes* Attems, 1930 from South and Southeast Asia. First, we determined the phylogenetic position of the genus *Digitipes* within the Order Scolopendromorpha (n=414) by assembling primary and published sequences (n=30) for two mtDNA markers (COI, 16S) and one nuclear marker (28S) using Maximum Likelihood and Bayesian inference. We further used single-locus and multi-locus coalescent-based species delimitation methods to identify putative species within the genus *Digitipes.* We then used three fossil calibrations to estimate divergence times in a Bayesian framework, and used the resulting time-calibrated phylogeny for biogeographic analysis in a likelihood framework (BioGeoBEARS). The genus *Digitipes* was monophyletic with strong support, with SEA lineages nested within the Indian clade and sister to the *D. barnabasi* species complex from the Western Ghats. *D. pruthii*, the Eastern Ghats species, was nested with the Western Ghats species clade. A single-locus mPTP-based species-delimitation method suggested the presence of 24 putative species, far exceeding the number of morphologically described species, indicating an underestimation of species diversity. Divergence time estimates suggest that *Digitipes* began diversifying around 126 mya (100-159 mya), affirming its Gondwanan origin. Time-stratified ancestral area reconstruction suggested that early vicariance, followed by jump dispersal and range expansion, shaped the distribution of the genus *Digitipes* in South and Southeast Asia. There was one dispersal event from India to Southeast Asia, following a transient land connection between them, around 50 mya, supporting the Out-of-India hypothesis. Additionally, three jump dispersal events and five range expansions explained diversification within peninsular India. Particularly, *D. pruthii* originated from a jump dispersal event from the Central Western Ghats to the Eastern Ghats around 37 mya. Our results highlight the importance of an integrative taxonomic framework to delineate hidden diversity and to obtain robust species hypotheses for testing biogeographic hypotheses.

## Introduction

Understanding the role of biogeographic processes in shaping current diversity and distribution patterns remains fundamental to ecology. The relative importance of vicariance and dispersal in determining current biodiversity patterns varies across regions and is influenced by their unique geoclimatic histories. For instance, Andean tectonics influenced the historical diversification of Neotropical biodiversity through dispersal and vicariance across subregions at different times (Antonelli et al., 2009; Luebert & Weigend, 2014; Meseguer et al., 2022). The breakup of the Gondwana landmass, subsequent plate collisions and associated climate history played a key role in South and Southeast Asian (SEA) biogeography (Lohman et al., 2014; Skeel et al., 2023). Understanding the biogeography of the peninsular Indian biota, particularly the roles of vicariance and dispersal, remains challenging due to the complex geo-climatic history. The Peninsular Indian Plate (PIP) of the Indian subcontinent was part of Gondwanaland 200 million years ago (mya) (Mani, 1974; Briggs, 2003). Approximately 120-90 mya, it separated from the supercontinent along with Madagascar, and PIP began drifting northward toward the Eurasian plate (Briggs, 2003; Ali & Aitchison, 2008). It continued its northward journey after separating from Madagascar and the Seychelles around 86-88.5 and 66 mya, respectively (Storey et al., 1995; Collier et al., 2008). Towards the end of the Cretaceous, around 66 mya, massive volcanic activity formed the Deccan Traps and coincided with mass extinctions and drastic climate change across PIP (Courtillot & Renne, 2003; Renne et al., 2013; Schoene et al., 2015; Keller et al., 2020). Post-volcanism, at the end of the Palaeocene and early Eocene (66-47 mya), dense perhumid rainforests were established throughout the PIP as it approached the equator (Prasad et al., 2009; Agnihotri et al., 2025; Parmar et al., 2025). This was followed by India’s collision with Eurasia around 40-35 mya, leading to biotic exchange between the landmasses (Ali & Aitchison, 2008; Klaus et al., 2016; Morley, 2018; Joshi et al., 2020). During its final collision with Eurasia towards the end of the Eocene (∼35 mya onwards), evidence suggests a more seasonal climate than the earlier perhumid one (Dupont-Nivet et al., 2007; Ali & Aitchison, 2008; Zachos et al., 2008; Clementz et al., 2011; Spicer, 2017; Morley, 2018). Although the exact timing of the collision remains unclear, studies show that, rather than a single collision event, multiple land bridges formed between India and Eurasia spanning approximately 55 mya to 20 mya (Ali & Aitchison, 2008; Najman et al., 2010; Hu et al., 2016; Klaus et al., 2016; Morley, 2018). Given the geo-climatic history of the PIP, it has been hypothesised that the PIP acted as a “biotic ferry” as these events leading up to the collision facilitated dispersal events across taxa in and out of the Indian subcontinent across similarly wet regions of SEA (Ali & Aitchison, 2008; Klaus et al., 2016; Morley, 2018). This is also known as the Out-of-India hypothesis, which posits that India is an ancestral area and that there was a dispersal event from India to SEA (Bossuyt & Milinkovitch, 2001; Datta-Roy & Praveen Karanth, 2009) (Fig. 1). Some of the older lineages found in the SEA have supported this hypothesis, e.g., centipede genus *Rhysida* (Joshi et al., 2020), Ochyroceratid spiders (Li et al., 2020), Gecarcinucidae crabs (Klaus et al., 2009), *Ichthyophis* (Gower et al., 2002) and Chikilidae caecilians (Kamei et al., 2012). In addition, the Burma Terrane hypothesis suggests that India was in contact with the Burma Plate c. 50 mya, and dispersal events have been documented among terrestrial lineages from India to SEA (Li et al., 2020; Bansal et al., 2022; Najman et al., 2022).

**Fig. 1.**
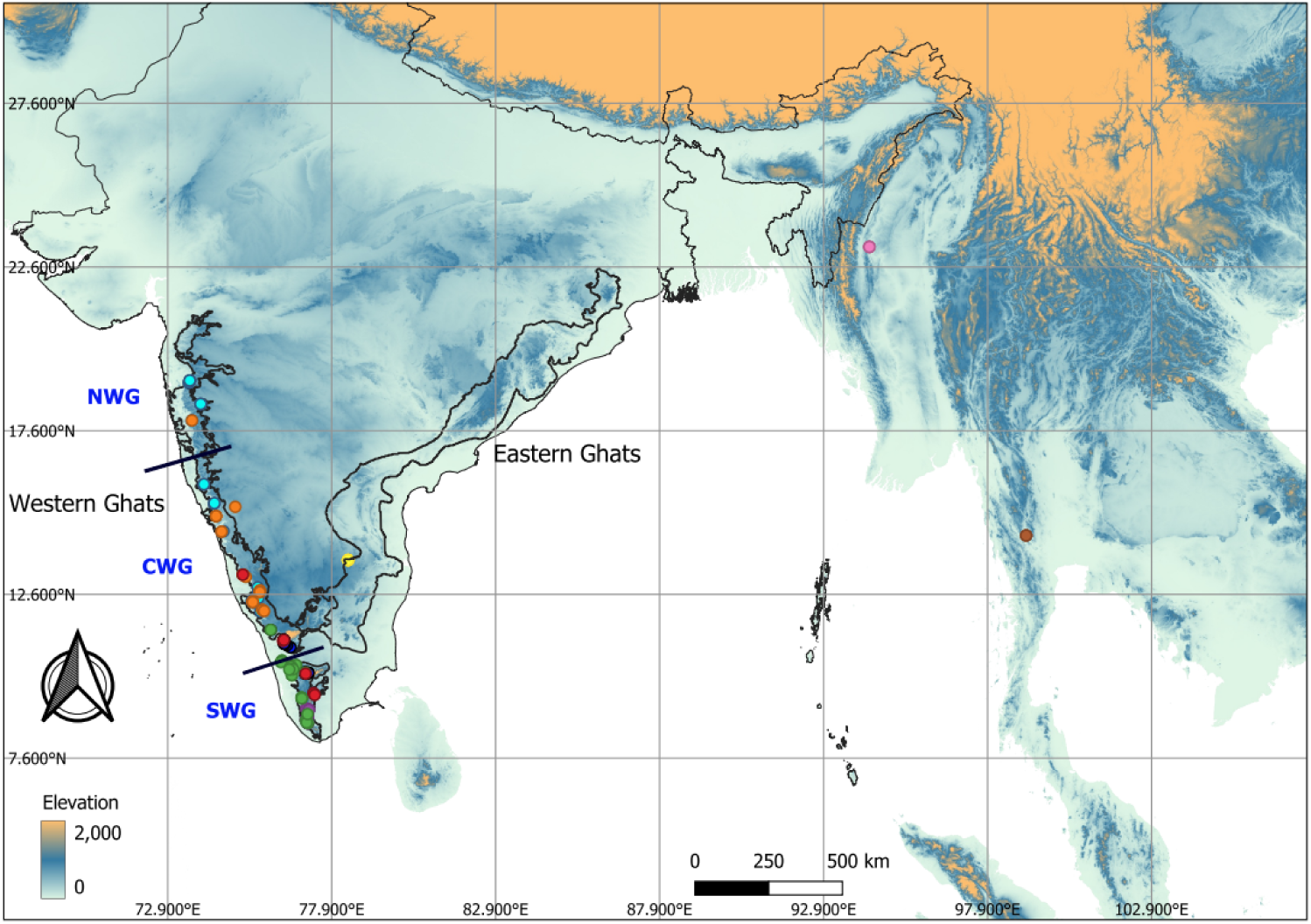
The centipede genus *Digitipes* taxon sampling from the Western and Eastern Ghats in peninsular India and Southeast Asia.

Furthermore, in situ diversification continued in PIP across lineages during the Cenozoic, leading to endemic radiations thought to have been facilitated by climatic shifts. Post-collision, the Himalayan uplift began at the end of the Eocene and intensified only in the mid-Miocene, coinciding with global warming during the mid-Miocene (16-11.5 mya) (Morley, 2018). The contraction of wet forests in PIP during the Late Miocene and Pliocene (11-2.5 mya) is potentially caused by subsequent regional drying and cooling, and there was also a simultaneous expansion of C4 grasslands (Morley, 2018). Therefore, peninsular India’s once-continuous wet forests are now restricted predominantly to the western slopes of the Western Ghats (WG) and to isolated wet forest pockets on the mountaintops of the Eastern Ghats (EG). However, our understanding of the relative roles played by climatic or geographic vicariance and dispersal among these lineages remains limited.

One of the ancient lineages among soil arthropods, the centipede genus *Digitipes* Attems 1930 (Order Scolopendromorpha), is mainly found in the wet forests of Africa in the Congo basin (three species), peninsular India (seven species) (Joshi & Edgecombe, 2013) and recently found in the wet forests of SEA (Siriwut et al., 2015), offering an opportunity to study both Out-of-India and diversification within PIP. A phylogenetic and biogeographic study reported that the genus *Digitipes* began diversifying in peninsular India during the Late Cretaceous, suggesting a Gondwanan origin (Joshi & Karanth, 2011, 2013). Therefore, we first tested predictions of the Out-of-India hypothesis to examine whether peninsular Indian species were ancestral to SEA species and whether the dispersal event occurred post-collision with Eurasia. We also rediscovered *D. pruthii,* an endemic species of mountaintops in the Eastern Ghats, 50 years after its description. With this additional species, we have sampled all described species of the genus *Digitipes* in peninsular India, a rarity among tropical soil arthropods.

Additionally, each species’ distributional range has been revised using species distribution models with primary field data. With nearly complete taxon sampling, revised distributional and molecular data for this ancient endemic group enabled us to examine how wet-forest fragmentation may have influenced *Digitipes* diversification in peninsular India. Specifically, we examined if the divergence between *D. pruthii* and the closest WG species corresponds with (1) cooling and aridification towards the end of the Eocene (∼34 mya), (2) more pronounced aridification during the Late Miocene (11–5 mya), or (3) Pleistocene climatic fluctuations (1.81– 0.01 mya). We also examined the relative roles of vicariance and dispersal in driving this divergence. We used molecular phylogenies to assess evolutionary relationships among species, species delimitation methods to delineate species boundaries, and a Bayesian approach to estimate divergence times. We then used time-stratified historical biogeographic analyses in a Maximum Likelihood framework to test the predictions of the Out-of-India hypothesis and the relative roles of climatic/geographic vicariance and dispersal within peninsular India (Figure 1).

## Methods

### Taxon Sampling and DNA Sequencing

A total of 31 new *Digitipes* specimens, including *D. pruthii*, were collected across the WG and EG (Figure 2, Table S1). The specimens were preserved in 70% ethanol. Morphological identification was carried out using published taxonomic keys, and DNA sequence data were generated for two mtDNA markers and one nuclear marker (Table S1). This was complemented by published DNA sequence data for *Digitipes,* representing seven species and 109 individuals across all three markers (Table S2). We used the manual phenol-chloroform-isoamyl alcohol method to extract DNA from tissue from 2-4 legs, depending on the individual’s size. Two mitochondrial markers (COI, 16S) and one nuclear marker (28S) were amplified using Polymerase Chain Reaction (PCR) following the protocols described in Joshi & Karanth, 2012 (Primer details and cycling conditions provided in Table S3) and subsequently sequenced using in-house Sanger sequencing at the CSIR-Centre for Cellular and Molecular Biology facility in Hyderabad. The final concatenated dataset had a total length of ∼1500 bp across 166 taxa, including outgroups.

**Fig 2.**
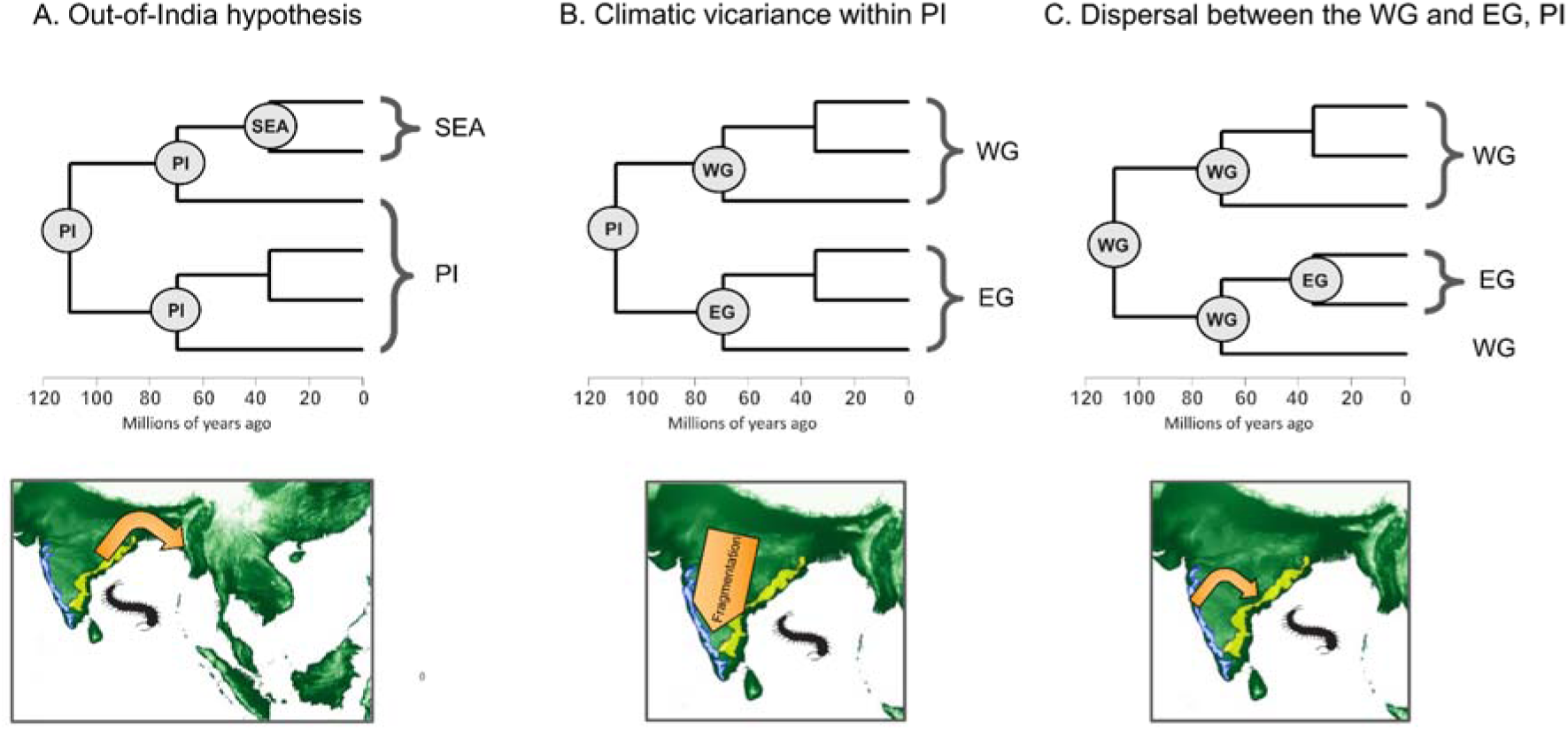
Phylogenetic predictions along with illustrated maps for the A) Out-of-India hypothesis, B) Climatic vicariance within peninsular India and C) Dispersal between the Western and Eastern Ghats, peninsular India

### Molecular Phylogenetic Analysis

DNA sequences for individual markers were aligned using the MUSCLE algorithm in MEGA 12 (Kumar et al., 2024). Maximum Likelihood and Bayesian approaches were used to reconstruct molecular phylogenetic trees for the genus *Digitipes*. First, to ascertain the phylogenetic position of the genus *Digitipes* within the subfamily Otostigminae, we assembled data for multiple genera and individuals across all three markers. We used *Scolopendra*, *Cormocephalus, and Asanada* from the sister family, Scolopendridae, as outgroups (n= 26).

PartitionFinder2 (Lanfear et al., 2016) was used to determine the optimal partitioning scheme and the best-fit models of sequence evolution for each partition based on the Bayesian Information Criterion (BIC) and a greedy search algorithm. Maximum likelihood (ML) analyses were conducted on the concatenated dataset using ModelFinder implemented in the IQ-TREE web server. The selected partitioning scheme consisted of five partitions: F81+F+G4 for COI codon position 1, TIM2+F+G4 for COI codon position 2, TNe+G4 for COI codon position 3, TVM+F+R5 for 16S, and TIM2e+G4 for 28S. ML tree reconstruction was performed by defining separate partitions for the three COI codon positions and one partition each for 16S and 28S. Branch support was assessed using the ultrafast bootstrap method with 1000 replicates.

Next, the best-fit models for each partition, as suggested by PartitionFinder2 from a set of only MrBayes models (Ronquist et al., 2012), were used to generate a Bayesian phylogenetic tree. The analysis was performed in MrBayes 3.2.7a (Ronquist et al., 2012) on a high performance computing cluster (HPC) for 500 million generations with 4 chains, 2 runs and a sampling frequency of 1000. The average standard deviation of split frequencies towards the end of the runs was below 0.01, thus indicating that both runs had converged. Tracer 1.7.2 (Rambaut et al., 2018) was used to assess when stationarity was achieved, and the first 10% of the trees were discarded as burn-in. Finally, a consensus tree was built from the remaining trees. Both ML and BI trees were visualised with FigTree v1.4.4 (Rambaut, 2018).

### Species delimitation and divergence time analysis

To generate a robust species hypothesis for the genus *Digitipes*, we applied three species-delimitation approaches. The two exploratory methods were mPTP (Multi-rate Poisson tree processes) (Kapli et al., 2017), a coalescent-based species delimitation method using single-locus sequence and tree data, and ASAP (Assemble Species by Automatic Partitioning) (Puillandre et al., 2021), a distance-based partitioning method. The third approach was StarBEAST2 (Bouckaert et al., 2014; Ogilvie et al., 2017), a validation method based on multi-locus data.

mPTP is a single-locus species-delimitation method that detects the minimum branch length from the alignment provided. The algorithm splits the input tree into sub-clades, fits different exponential distributions (i.e., multi-or rate of branch lengths) for each putative species-clade, and selects the overall best-fit splitting scheme as the best model for species delimitation.

Further, it uses the MCMC method to sample all possible spatial delimitations, providing support values for the resulting delimitation scheme. IQTREE (Nguyen et al., 2015) was used to build a maximum-likelihood phylogenetic tree from 140 individuals, with a combined COI and 16S dataset treated as a single locus for *Digitipes,* using *Rhysida immarginata* as the outgroup. The rooted, binary likelihood tree and the aligned FASTA sequences were used for mPTP analysis. First, a minimum branch length of 0.0008577573 was evaluated using the tree and FASTA files, which the algorithm used to distinguish interspecific divergences from intraspecific coalescences. Next, MCMC delimitation analysis was performed, starting from the null model that treats the entire tree as a single species. Four independent MCMC runs were performed for 10 million generations, sampling every 1000 generations and discarding the first 1 million as burn-in. A generation-versus-log-likelihood plot was utilised to assess the convergence of the individual runs. The results of this analysis were utilised to define taxon sets for the multilocus StarBEAST2 analysis (Table S4).

ASAP was used to perform species delimitation on a subset of the COI dataset for *Digitipes* (117 sequences, 530 bp) (Table S5). ASAP partitions sequences into putative species based on pairwise genetic distances, with lower ASAP scores indicating better partitioning. We evaluated the 10 best ASAP partitions, ranked by ASAP score, and reported the best-ranked partition. (Table S6). For each cluster, we assessed cluster uniformity and cluster size to indicate the clarity and reliability of ASAP-assigned clusters (Table S5).

To obtain a species tree with divergence date estimates, StarBEAST2 analysis was implemented in BEAST 2.6.7 (Bouckaert et al., 2014), utilising a multi-species coalescent framework under a relaxed exponential clock model. For this analysis, we used a larger concatenated three-marker dataset (COI, 16S, and 28S) comprising 414 taxa and multiple outgroups from different centipede orders (Supplementary material 1). Three fossil-based calibrations with log-normal distribution priors and four secondary calibrations with uniform priors were added. Additionally, monophyly constraints were imposed on nodes to ensure topological consistency with the species tree and the trees obtained by IQ-TREE and MrBayes. Secondary calibrations were set according to a published, dated phylogeny of the scolopendrid centipede genus *Rhysida* (Joshi et al., 2020). The details of all fossil and secondary calibrations, their lower and upper age bounds, and references are provided in Table S7. First, PartitionFinder2 was used to select the best-fitting partition scheme and model of sequence evolution from a set of StarBEAST2 models only. A three-partition scheme was selected, with the GTR + I model of sequence evolution for COI, 16S and 28S. Thus, the substitution models were linked across loci, and the GTR model was selected for all three partitions. Clock models and gene trees were linked for mitochondrial genes (COI, 16S), which were treated as a single locus separate from the nuclear gene (28S). Next, we reconciled morphology, geographic distribution, mPTP-based species delimitation, and existing phylogenetic relationships to define new species hypotheses for the genus *Digtipes* (Table S8). Gene ploidy was set to haploid for mitochondrial genes (COI, 16S) and diploid for the nuclear gene (28S). The Yule speciation process was selected prior, and “Linear with constant root populations” was selected as the population model.

The analysis was run on HPC for 500 million generations, with a sampling frequency of 5,000. The convergence of priors was visualised in Tracer 1.7.2 (Rambaut et al., 2018), and their Effective Sample Size (ESS) values were confirmed to be at least 200. The first 10% of the trees were discarded as burn-in. The species trees obtained from this analysis were resampled with 10,000 frequency and 25% burn-in in LogCombiner (Drummond & Rambaut, 2007). Finally, TreeAnnotator (Drummond & Rambaut, 2007) was used to get a Maximum Clade Credibility (MCC) tree, which was visualised using FigTree v1.4.4.

#### Historical biogeography analyses

We performed time-stratified ancestral area reconstruction on a time-calibrated *Digitipes* phylogeny and its sister group, using the R (R Core Team, 2024) package BioGeoBEARS v.1.1.3 (Matzke, 2013a), to infer ancestral ranges and compare models of range evolution. To account for the biogeographic history within the species complexes and variation in geographic ranges among them, we performed this analysis based on the 15 distinct clades. A total of six models were implemented in a maximum likelihood framework: Dispersal-Extinction Cladogenesis (DEC) (Ree & Smith, 2008), likelihood versions of Dispersal-Vicariance (DIVAlike) (Ronquist, 1997; Matzke, 2013b) and Bayesian Analysis of Biogeography (BAYAREAlike) (Landis et al., 2013; Matzke, 2013b), along with DEC+J, DIVAlike+J, BAYAREAlike+J models, that include an additional parameter +J representing “jumps” or a founder-speciation event (j) (Matzke, 2013b).

These models were implemented in a series of time-stratified analyses to compare run results with the following inputs and settings: 1) A1 run constrained with the ‘area function’ in time strata but no constraint on between area dispersal probabilities and 2) A2 run constrained with the ‘area function’ in time strata and constrained between area dispersal probabilities (details on ‘area function’ involving manual modification of states provided in Supplementary material 3, point 2.). Analyses without dispersal constraints used dispersal matrices with equal probabilities among all areas (i.e., 1).

We defined five different subregions as areas within the Oriental realm: Mainland Southeast Asia (A), Eastern Ghats (E), and the Western Ghats, which were further divided into Northern Western Ghats (N - NWG), Central Western Ghats (C - CWG), and Southern Western Ghats (S - SWG) based on geoclimatic barriers and phytogeography. We performed time-stratified biogeographic analysis using the following time points based on geoclimatic events of the Indian subcontinent: 150 mya, chosen as a time point older than the estimated crown age of *Digitipes* (148 my), 65 mya as the time of Deccan volcanism event, 55 mya as the time of first ever transient land connection with Asia during India’s northward journey, and 20 mya as the beginning of climatic shifts that caused aridification, and thus may have influenced wet-forest distribution and diversification of *Digitipes* within the peninsular region. (Table S9)

The maximum range size was set to 4. In both A1 and A2 analyses, non-adjacent ranges were manually disallowed to define the allowed areas for each time stratum. Southeast Asia (A) was considered non-adjacent to the remaining areas post 20 mya, due to the drier Indo-Gangetic plains acting as a dispersal barrier for wet forest-adapted taxa (Morley, 2018) (Supplementary material 3, point 2).

For each time stratum, the probability for contiguous areas was set to 1; adjacent areas with moderate connectivity but a barrier were set to 0.75; moderately separated areas with intermediate connections, or separated by another area/two or more landmasses, were set to 0.5; well-separated areas by water/large terrestrial barrier were set to 0.00001. In the A2 analysis, dispersal probability between SEA and other subregions was increased from 0.00001 to 0.5, as it corresponded to India’s multiple contacts with SEA during the 55-20 mya time strata (Table S9). PIP-SEA dispersal probability was again decreased post 20 mya, as during this time the drier Indo-Gangetic plains could have acted as a dispersal barrier for wet forest taxa, reducing the Out-of-India dispersals. Moreover, dispersal probabilities between areas within PIP post 20 mya were reduced because of reduced connectivity among these regions, resulting from wet-zone fragmentation driven by aridification. The details of the dispersal matrices, including all dispersal probabilities for each time stratum in the A2 analysis, are provided in Table S10.

AICc (Akaike Information Criterion corrected for small sample sizes) was used to choose the best-fitting model for our data, and LRT (Likelihood Ratio Tests) were used to compare nested models (DEC and DEC+J, DIVAlike and DIVAlike+J, BAYAREAlike and BAYAREAlike+J).

## Results

### Phylogenetic relationships and species delimitation within the genus *Digitipes*

The Maximum Likelihood and Bayesian approaches recovered similar topologies for the Otostigminae subfamily, in which the genus *Digitipes* was monophyletic with high support (Fig. 3). There were two clades within *Digitipes*: one consisted of WG and EG endemic species (Clade A), and the other (Clade B) consisted of SEA species (*D.* sp. and *D. kalewaensis*) along with the *D. barnabasi* species complex from the WG. The species delimitation methods, mPTP and ASAP, suggested the presence of 24 and 35 distinct putative species, respectively. We reconciled evidence from species delimitation analyses, monophyly, geographic distribution, and morphology to propose a new, robust, albeit conservative species hypothesis for the genus *Digitipes* with 10 species (Figure 3). *D. coonoorensis* 1 and 2, *D. jonesii*, *D. pruthii*, *D. nudus, and D. jangii* were part of Clade A. Among these, *D. coonoorensis* was inferred to be a species complex comprising two distinct putative species, which were allopatric in distribution. Additionally, a distinct species, *D.* sp. A, was found in the southern and central WG. We also recovered multiple distinct units within *D. jonesii* and *D. jangii*; however, they had overlapping geographic distributions and lacked diagnostic characters, so we treated them as a single species in this study. Clade B included two species from Southeast Asia, *D.* sp. and *D. kalewaensis,* which were sister to the *D. barnabasi* species complex. Within *D. barnabasi,* four distinct units were recovered, which were largely allopatric in distribution, though they lacked diagnostic morphological characters and are currently treated as a single species.

**Fig 3.**
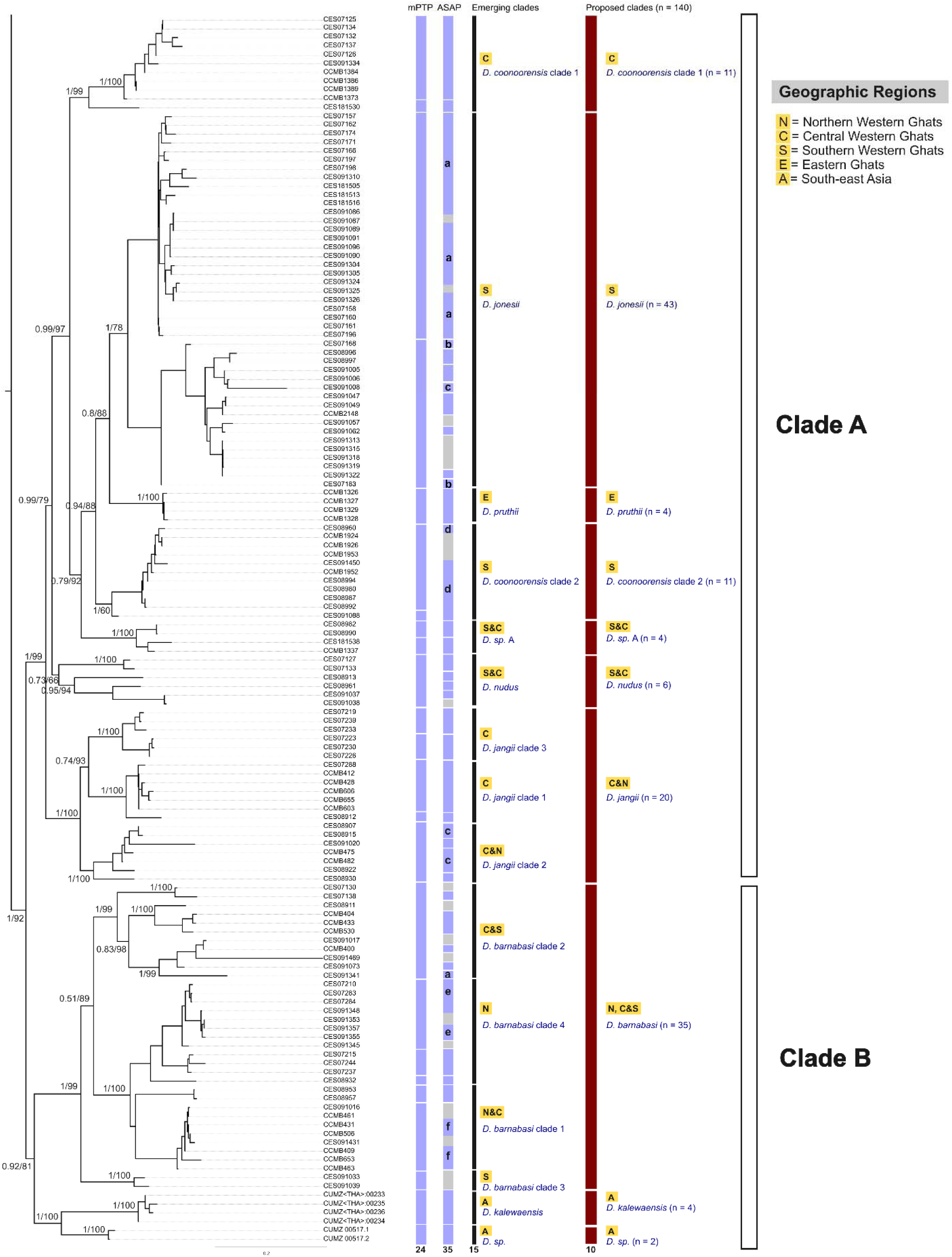
A Bayesian tree and species delimitation results (mPTP and ASAP) for the genus *Digitipes* with posterior probability and bootstrap support values.

### Divergence time estimation

The genus *Digitipes* began diversifying around 126 mya (95% HPD: 100-159 mya). Clade A, consisting of *D. coonoorensis* 1 and 2, *D. jonesii*, *D. pruthii*, *D. nudus,* the *D. jangii* species complex and the new *D.* sp. A., originated around 104 mya (95% HPD: 80-133 mya). Clade B, consisting of *the D. barnabasi* species complex, the Southeast Asian species *D.* sp., and *D. kalewaensis,* originated around 107 mya (95% HPD: 78-141 mya). Within Clade A, *D. nudus* diverged from *D. jangii* around 89 mya (95% HPD: 65-115 mya), and the *D. jangii* species complex began diversifying around 75 mya (95% HPD: 47-104 mya). The new *D.* sp. A diverged from *D. jonesii* around 54 mya (95% HPD: 23-85 mya), and *D. coonorensis* 2 diverged from *D. pruthii* and *D. coonoorenis* 1 around 58 mya (95% HPD: 33-79 Mya). The EG species *D. pruthii* and *D. coonoorensis* diverged around 37 mya (95% HPD: 8-67 mya). Within Clade B, the *D. barnabasi* species complex began diversifying around 92 mya (95% HPD: 63-124 mya), while *D. sp.* and *D. kalewaensis* diverged around 48 mya (95% HPD: 3-94 mya) (Figure 4).

**Fig 4.**
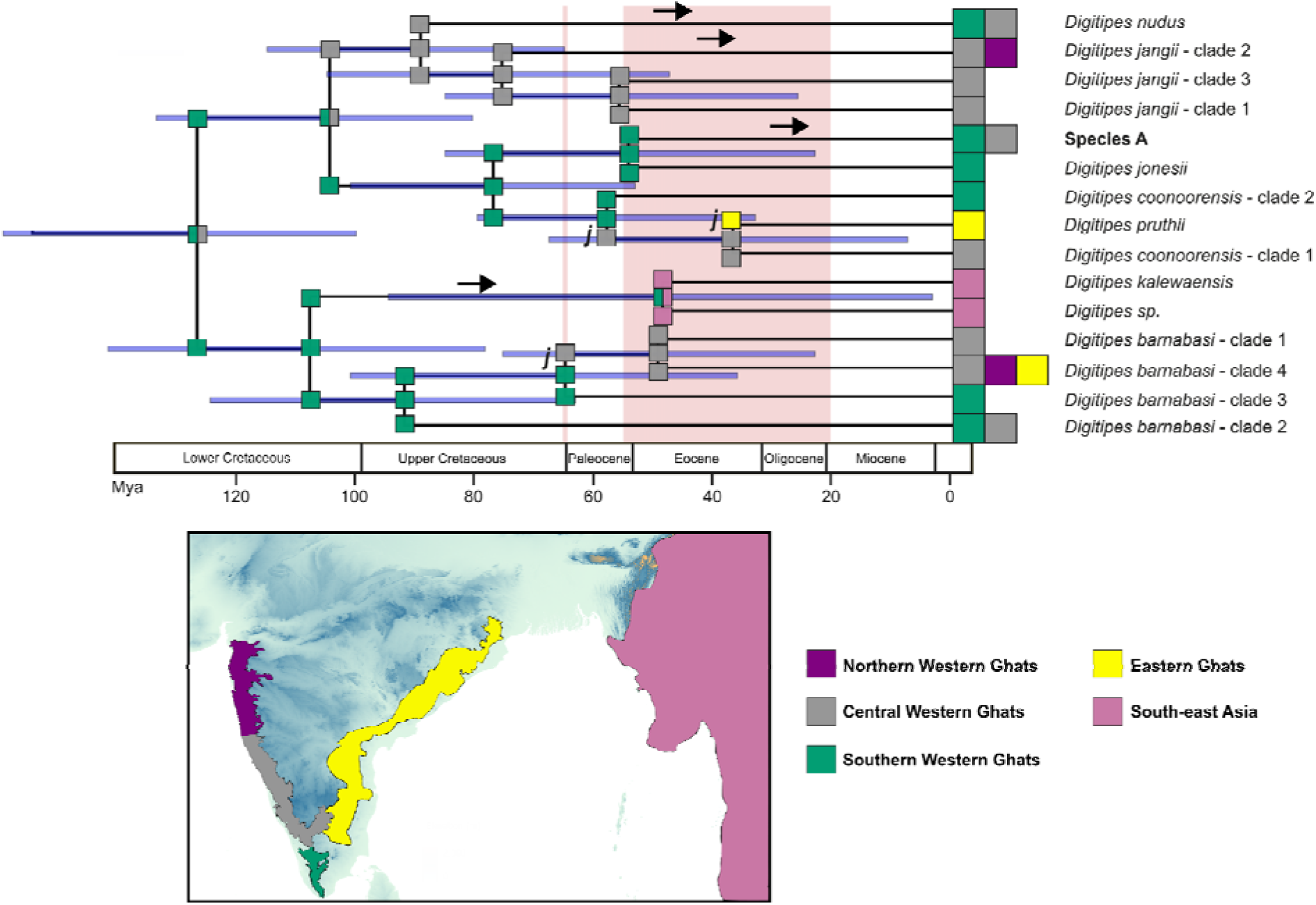
A time-tree for the genus *Digitipes* with reconstructed ancestral areas (BioGEOBEAR: DEC+J); ancestral areas are shown at each node, dispersal events are shown in black arrows, and a jump dispersal event is indicated by j.

### Ancestral range estimation

Among the historical biogeography models, the DEC+J model, in which dispersal probability from PIP to SEA was higher during the 55-20 mya time interval (A2 scenario), had the lowest AICc, with a log-likelihood of -38.27 (Table 1). The historical biogeographic reconstruction indicated that the ancestral range of the genus *Digitipes* was in SWG-CWG, followed by vicariance, with Southern and Central WG remaining ancestral areas for Clade A and SWG for Clade B. Another vicariant event around 104 mya, in which CWG became an ancestral area for a clade comprising *D. nudus* and *D. jangii* in CWG, and SWG for the remaining species in Clade A. A jump dispersal event from SWG to CWG led to *D. coonoorensis* 2 around 58 mya, and another jump dispersal leading to *D. pruthii* occurred from Central WG to EG post 37 mya. The third jump dispersal event occurred in the *D. barnabasi* species complex, from SWG to CWG around 65 mya, followed by range expansions in the NWG and EG.

**Table 1.** BiogeoBEARS model comparison results for time-stratified analyses under hypotheses A2 and A1. Time-stratified analyses were run using hypothesis A2 (Variable dispersal matrix with time-stratified area-function constraints) and hypothesis A1 (Equal dispersal matrix with time-stratified area-function constraints). For each model, log-likelihood (LnL), number of parameters, estimated dispersal (d), extinction (e), and jump dispersal (j) parameters, Akaike Information Criterion corrected for small sample sizes (AICc), ΔAICc, and AICc weights are reported. The highlighted rows refer to the two models with the lowest AICc scores.

| Model | LnL | numparams | d | e | j | AICc | Delta_AICc | AICc_weight |
| --- | --- | --- | --- | --- | --- | --- | --- | --- |
| **Constrained with the 'area function' in time strata and constrained between-area dispersal probabilities (A2)** |  |  |  |  |  |  |  |  |
| DEC | -40.61 | 2 | 0.0035 | 0.0017 | 0 | 86.21 | 1.48 | 0.183 |
| DEC+J | -38.27 | 3 | 0.0026 | 0.0011 | 0.12 | 84.73 | 0 | 0.385 |
| DIVALIKE | -41.11 | 2 | 0.0041 | 0.0014 | 0 | 87.22 | 2.49 | 0.111 |
| DIVALIKE+J | -39.18 | 3 | 0.003 | 0.0011 | 0.11 | 86.55 | 1.82 | 0.155 |
| BAYAREALIKE | -42.65 | 2 | 0.0021 | 0.0075 | 0 | 90.3 | 5.57 | 0.024 |
| BAYAREALIKE+J | -40.97 | 3 | 0.0025 | 0.0013 | 0.14 | 90.13 | 5.4 | 0.026 |
| **Constrained with the 'area function' in time strata but no constraint on between-area dispersal probabilities (A1)** |  |  |  |  |  |  |  |  |
| DEC | -56.5 | 2 | 0.01 | 0.01 | 0 | 118 | 33.27 | 2.29E-08 |
| DEC+J | -56.5 | 3 | 0.01 | 0.01 | 0.0001 | 121.2 | 36.47 | 4.63E-09 |
| DIVALIKE | -42.84 | 2 | 0.0027 | 0.0014 | 0 | 90.69 | 5.96 | 0.020 |
| DIVALIKE+J | -39.96 | 3 | 0.0019 | 0.001 | 0.1 | 88.11 | 3.38 | 0.071 |
| BAYAREALIKE | -43.4 | 2 | 0.0014 | 0.0073 | 0 | 91.8 | 7.07 | 0.011 |
| BAYAREALIKE+J | -41.51 | 3 | 0.0016 | 0.0012 | 0.12 | 91.2 | 6.47 | 0.015 |

See Appendix for detailed descriptions of A1 and A2 scenarios, the dispersal multipliers applied in each case, and the R implementation of the 'area function'.

**Table 2.** Likelihood ratio test between three standard biogeography models (DEC, DIVALIKE, BAYAREA), each nested within their “+J” counterpart (for Hypothesis A2). Here, the “+J” event is helpful in explaining the data in the DEC model (p < 0.05).

| Alternative model | Null model | LnLalt | LnLnull | DFalt | DFnull | DF | Dstatistic | p-value | test | tail |
| --- | --- | --- | --- | --- | --- | --- | --- | --- | --- | --- |
| DEC+J | DEC | -38.27 | -40.61 | 3 | 2 | 1 | 4.68 | 0.031 | chi-squared | one-tailed |
| DIVALIKE+J | DIVALIKE | -39.18 | -41.11 | 3 | 2 | 1 | 3.86 | 0.049 | chi-squared | one-tailed |
| BAYAREALIKE+J | BAYAREALIKE | -40.97 | -42.65 | 3 | 2 | 1 | 3.36 | 0.067 | chi-squared | one-tailed |

Additionally, a dispersal event from SWG to the SEA occurred around 48 mya, making SWG-SEA the ancestral area for two SEA species. Overall, there have been two vicariance events, three jump dispersals, and seven range expansions across the biogeographic history of the genus *Digitipes* (Figure 4).

## Discussion

### Systematics of *Digitipes*

The molecular phylogeny confirms that *Digitipes* is a well-supported monophyletic genus within Otostigminae, comprising two major clades: one restricted to the Western and Eastern Ghats of peninsular India, and another where SEA species were nested within Indian lineages. *D. pruthii* was rediscovered after ∼ 40 years since its description (Jangi & Dass, 1984). Species delimitation analyses reveal a much higher putative species diversity (24 in mPTP, 35 in ASAP) than previously recognised morphologically (8 morphospecies), suggesting significant cryptic diversity within *Digitipes* (Figure 3). No diagnostic morphological characters could be identified to distinguish the putative species. This highlights the need for an integrative taxonomy combining molecular, morphological, and geographic data to understand and delineate species boundaries in soil arthropods. The findings also highlight complex patterns of lineage diversification in the Western Ghats, a known biodiversity hotspot, with multiple species complexes.

### Support for Out of India Hypothesis

The phylogenetic nested position of Southeast Asian lineages within the Indian clade, together with a transient Eocene land connection (∼50 mya), supports a single dispersal event from PIP into Southeast Asia, consistent with the Out-of-India hypothesis. This pattern aligns with geoclimatic evidence of multiple biotic exchanges before the final India-Eurasia collision (Ali & Aitchison, 2008; Najman et al., 2010; Hu et al., 2016; Klaus et al., 2016; Morley, 2018), and supports the ‘biotic ferry’ hypothesis, wherein wet-forest taxa such as *Digitipes* expanded their ranges into Southeast Asia, with further dispersal later constrained by climatic shifts and geographic barriers such as the Indo-Gangetic plains (Morley, 2018).

Divergence-time estimates suggest the origin of the genus *Digitipes* was ∼126 Mya (95% HPD: 100-159 mya), overlapping with the separation of peninsular India from Gondwana (∼120-90 mya), consistent with a Gondwanan vicariant origin for the genus (Joshi & Karanth, 2011, 2013). Following this ancient origin, *Digitipes* diversified extensively within peninsular India before a subsequent dispersal event connected it to Southeast Asia.

### Implications for Peninsular Indian Biogeography

Within peninsular India, the diversification of *Digitipes* reflects the region’s complex geo-climatic history and habitat fragmentation. The split between Eastern Ghats and Western Ghats species, and the subsequent range expansions, correspond to known climatic transitions, including post-Eocene aridification and Late Miocene drying episodes that fragmented once-continuous wet forests (Morley, 2018). The dispersal of *D. pruthii* from the Central Western Ghats to the Eastern Ghats around 37 mya illustrates that these isolated wet forest patches were once connected, facilitating dispersal; and dispersal from one wet forest patch to another implies a role for niche conservatism in shaping *D. pruthii’*s distribution.

Reduced dispersal probabilities after 20 mya, especially between subregions within peninsular India, indicate that climatic drying and habitat isolation have shaped current distribution patterns by promoting vicariance. Overall, the findings emphasise the importance of historical climate and geological events in generating endemic biodiversity through both in-situ diversification and limited dispersal within the peninsular wet forests.

This long evolutionary history and persistence of the genus *Digitipes* through major geological and climatic events, such as the breakup of Gondwana and the northward drift of the Indian Plate, suggest resilience and evolutionary stability within wet-forest ecosystems. The splits between the WG and EG clades and between the SEA and Indian clades highlight the role of geographic isolation within peninsular India and between India and SEA in promoting lineage diversification. These patterns imply that geological changes and wet-forest fragmentation created ecological and evolutionary barriers early on, contributing to the endemism. The multiple dispersals and range expansions within Western Ghats clades before 20 mya, followed by reduced dispersal probabilities, point to a shift from widespread connectivity to increasing habitat fragmentation driven by Late Miocene-Pliocene aridification. This transition likely led to speciation by isolating populations in wet forest patches, emphasising climate as a major driver of present species distributions. The detection of far more putative species than those morphologically described, including species complexes within well-known taxa, indicates extensive hidden diversification, potentially driven by geoclimatic factors yet to be explored. Together, these illustrate how ancient geological events, periodic land connectivity, climatic fluctuations, and habitat fragmentation have interacted to shape the diversification and distribution of *Digitipes*. They reveal a dynamic biogeographic history with ongoing vicariance and dispersal, contributing to both regional endemism in peninsular India and biotic exchange with Southeast Asia.

This study provides a comprehensive framework that clarifies the systematics of the genus *Digitipes* and highlights the Indian subcontinent’s pivotal role in facilitating ancient biotic interchange and the diversification of wet-forest arthropods. It also highlights how integrating time-calibrated molecular phylogenies with data from past geological changes advances understanding of tropical biodiversity, its origins, and the evolutionary processes shaping species distributions.

## Author contributions

J.J. and P.D. conceptualised the study; P.D. collected the data and conducted the analyses along with J.J. and P.R.; P.D. and J.J. wrote the manuscript. All authors approved the final version.

## Acknowledgements

We thank Dr Bharti D. K., Dr Mihir Kulkarni, Mr Nehal Gurung, Ms Maya Manivannan, Ms Aditi Sinha, Mr Abhishek Gopal, Mr Sudhanshu Kumar, and Mr S.S. Dhanush for valuable discussions throughout the course of this study, assistance in fieldwork, and suggestions on the analyses. We thank the DNA sequencing facility at CSIR-Center for Cellular and Molecular Biology, Hyderabad, for the sequences generated and utilised in this study. We also thank the state forest departments of Andhra Pradesh, Kerala, and Karnataka for providing the necessary permits and assistance during fieldwork. This work was supported by the DBT/Wellcome Trust India Alliance Intermediate Fellowship [grant number IA/I/20/1/504919] and by the Council of Scientific & Industrial Research (CSIR), Government of India [grant number MLP0156], both awarded to JJ.

## Data Availability Statement

The DNA sequences generated in this study are deposited in GenBank (NCBI), and geographic distribution data are mentioned in the supplementary material 1.

## Supplementary material

Supplementary 1: List of taxa used in the phylogenetic analyses with their GenBank accession numbers and distribution.

Supplementary 2 (point 1) provides details of the laboratory protocol for DNA extraction and PCR, including primer details and cycling conditions. It also includes details on species delimitation, divergence time estimation, and historical biogeography analyses of the genus *Digitipes*.

