## Supplementary material 2 for "Biogeography and cryptic diversity of the ancient centipede genus *Digitipes* (Scolopendromorpha) in South and Southeast Asia"

### Supplementary Tables

S1. *Digitipes* Sampling in Peninsular India in this study

S2. Sequences used for *Digitipes* molecular phylogenetic analysis

S3. Primer details and cycling conditions

S4. *Digitipes* Species Delimitation (mPTP)

S5. ASAP analysis using the COI sequences of *Digitipes*

S6. 10 best ASAP ranked partitions for the *Digitipes* COI dataset

S7. Fossil Calibration for Divergence Time Estimation with starBEAST

S8. Species delimitation (mPTP) support and geography to determine emerging clades in *Digitipes*

S9. BioGeoBEARS analysis - Area Codes, Time Periods and Dispersal Multiplier Assignment

S10. BioGeoBEARS analysis - Dispersal matrices

#### S1. *Digitipes* Sampling in Peninsular India (Eastern and Western Ghats; n = 31) in this study

| **Specimen** | **Date** | **Species** | **Location** | **Latitude** | **Longitude** | **Elevation** | **DNA ng/ul** | **COI** | **16S rDNA** | **28S rDNA** |
| --- | --- | --- | --- | --- | --- | --- | --- | --- | --- | --- |
| CCMB400 | 14/08/21 | *Digitipes barnabasi* | Talacauvery WLS, Karnataka, India | 12.39842 | 75.45979 | 1013 | 5.9 | + | + | + |
| CCMB404 | 14/08/21 | *Digitipes barnabasi* | Talacauvery WLS, Karnataka, India | 12.39755 | 75.47372 | 1252 | 2.2 | + | + | + |
| CCMB409 | 15/08/21 | *Digitipes barnabasi* | Talacauvery WLS, Karnataka, India | 12.38568 | 75.49468 | 1229 | 15.1 | + | + | + |
| CCMB412 | 15/08/21 | *Digitipes jangii* | Talacauvery WLS, Karnataka, India | 12.36272 | 75.49148 | 1021 | 4.3 | + | + | + |
| CCMB428 | 15/08/21 | *Digitipes jangii* | Talacauvery WLS, Karnataka, India | 12.36272 | 75.49148 | 1021 | 13.7 | + | + | + |
| CCMB431 | 15/08/21 | *Digitipes barnabasi* | Talacauvery WLS, Karnataka, India | 12.3611 | 75.49017 | 981 | 8.6 | + | + | + |
| CCMB433 | 15/08/21 | *Digitipes barnabasi* | Talacauvery WLS, Karnataka, India | 12.38568 | 75.49468 | 1229 | 51.6 | + | + | + |
| CCMB461 | 17/08/21 | *Digitipes barnabasi* | Talacauvery WLS, Karnataka, India | 12.39995 | 75.53957 | 912 | 30.4 | + | + | + |
| CCMB463 | 17/08/21 | *Digitipes barnabasi* | Talacauvery WLS, Karnataka, India | 12.38893 | 75.49138 | 1211 | 23.4 | + | + | + |
| CCMB475 | 18/08/21 | *Digitipes jangii* | Pushpagiri WLS, Karnataka, India | 12.66079 | 75.70181 | 1096 | 10.4 | + | + | + |
| CCMB482 | 18/08/21 | *Digitipes jangii* | Pushpagiri WLS, Karnataka, India | 12.66739 | 75.70652 | 1066 | 18.6 | + | + | + |
| CCMB506 | 19/08/21 | *Digitipes barnabasi* | Pushpagiri WLS/Subramanya, Karnataka, India | 12.63081 | 75.64589 | 190 | 18.2 | + | + | + |
| CCMB530 | 20/08/21 | *Digitipes barnabasi* | Pushpagiri WLS/Galibeedu, Karnataka, India | 12.49116 | 75.66397 | 1002 | 6 | + | + | + |
| CCMB603 | 22/08/21 | *Digitipes jangii* | Makutta Reserve Forest, Karnataka, India | 12.12503 | 75.78373 | 700 | 8.4 | + | + | + |
| CCMB606 | 22/08/21 | *Digitipes jangii* | Makutta Reserve Forest, Karnataka, India | 12.12418 | 75.78588 | 635 | 8 | + | + | + |
| CCMB653 | 23/08/21 | *Digitipes barnabasi* | Makutta Reserve Forest, Kokka, Karnataka, India | 12.07755 | 75.72576 | 730 | 15.5 | + | + | + |
| CCMB655 | 23/08/21 | *Digitipes jangii* | Makutta Reserve Forest, Kokka, Karnataka, India | 12.09049 | 75.83879 | 799 | 14.9 | + | + | + |
| CCMB1326 | 20/10/21 | *Digitipes pruthii* | Horsley Hills, Andhra Pradesh, India | 13.64632 | 78.40147 | 1855 | 17 | + | + | + |
| CCMB1327 | 20/10/21 | *Digitipes pruthii* | Horsley Hills, Andhra Pradesh, India | 13.64632 | 78.40147 | 1855 | 12.1 | + | + | + |
| CCMB1328 | 20/10/21 | *Digitipes pruthii* | Horsley Hills, Andhra Pradesh, India | 13.64632 | 78.40147 | 1855 | 13.5 | + | + | + |
| CCMB1329 | 20/10/21 | *Digitipes pruthii* | Horsley Hills, Andhra Pradesh, India | 13.64632 | 78.40147 | 1855 | 4.8 | + | + | + |
| CCMB1337 | 18/06/22 | *Digitipes coonoorensis* | Silent valley national Park, Kerala, India | 11.0477 | 76.5367 | 526 | 56.3 | + | + | + |
| CCMB1373 | 19/06/22 | *Digitipes coonoorensis* | Silent valley national Park, Kerala, India | 11.0821 | 76.4774 | 866 | 36.5 | + | + | + |
| CCMB1384 | 21/06/22 | *Digitipes coonoorensis* | Silent valley national Park, Kerala, India | 11.0944 | 76.4471 | 982 | 33 | + | + | + |
| CCMB1386 | 21/06/22 | *Digitipes coonoorensis* | Silent valley national Park, Kerala, India | 11.0944 | 76.4471 | 982 | 25.1 | + | + | + |
| CCMB1389 | 21/06/22 | *Digitipes coonoorensis* | Silent valley national Park, Kerala, India | 11.0944 | 76.4471 | 957 | 28.5 | + | + | + |
| CCMB1924 | 22/08/22 | *Digitipes coonoorensis* | Anamudi Shola National Park, Kerala, India | 10.19954 | 77.1902 | 1733 | 48.8 | + | + | + |
| CCMB1926 | 22/08/22 | *Digitipes coonoorensis* | Anamudi Shola National Park, Kerala, India | 10.19954 | 77.1902 | 1733 | 26.6 | + | + | + |
| CCMB1952 | 23/08/22 | *Digitipes coonoorensis* | Anamudi Shola National Park, Kerala, India | 10.19883 | 77.19114 | 1767 | 19 | + | + | + |
| CCMB1953 | 23/08/22 | *Digitipes coonoorensis* | Anamudi Shola National Park, Kerala, India | 10.19883 | 77.19114 | 1767 | 32.2 | + | + | + |
| CCMB2148 | 23/08/22 | *Digitipes chhotanii* | Peechi Wildlife sanctuary, Kerala, India | 10.58805 | 76.33684 | 503 |  | + | + | + |

#### S2. Sequences used for molecular phylogenetic analysis of the genus *Digitipes* (n = 109)

| **Voucher No.** | **Tribe** | **Species** | **COI** | **16S rDNA** | **28S rDNA** | **Location** | **Reference** |
| --- | --- | --- | --- | --- | --- | --- | --- |
| CES07106 | Scolopendrini | *Scolopendra morsitans* | JN004006 | JN003895 | JN003951 | Ramanagar, Ramanagara district, Karnataka, India | Joshi and Karanth 2011 |
| CES07125 | Otostigmini | *Digitipes coonoorensis* | JN004032 | JN003921 | JN003976 | Silent Valley National Park, Kerala, India | Joshi and Karanth 2011 |
| CES07126 | Otostigmini | *Digitipes coonoorensis* | JN004033 | JN003922 | JN003977 | Silent Valley National Park, Kerala, India | Joshi and Karanth 2011 |
| CES07127 | Otostigmini | *Digitipes nudus* | JN004034 | JN003923 | JN003978 | Silent Valley National Park, Kerala, India | Joshi and Karanth 2011 |
| CES07130 | Otostigmini | *Digitipes barnabasi* | MK273205 | MK273311 | MK273433 | Silent Valley National Park, Kerala, India | Joshi and Edgecombe 2018 |
| CES07132 | Otostigmini | *Digitipes coonoorensis* | JN004035 | JN003924 | JN003979 | Silent Valley National Park, Kerala, India | Joshi and Karanth 2011 |
| CES07133 | Otostigmini | *Digitipes nudus* | JN004036 | JN003925 | JN003980 | Silent Valley National Park, Kerala, India | Joshi and Karanth 2011 |
| CES07134 | Otostigmini | *Digitipes coonoorensis* | JN004037 | JN003926 | JN003981 | Silent Valley National Park, Kerala, India | Joshi and Karanth 2011 |
| CES07137 | Otostigmini | *Digitipes coonoorensis* | JN004038 | JN003927 | JN003982 | Silent Valley National Park, Kerala, India | Joshi and Karanth 2011 |
| CES07138 | Otostigmini | *Digitipes barnabasi* | JX531697 | JX531827 | JX531776 | Silent Valley National Park, Kerala, India | Joshi and Karanth 2011 |
| CES07156 | Otostigmini | *Rhysida longipes* | MK273208 | MK273315 | MK273436 | Chatancode, Vidura, Kerala, India | Joshi and Edgecombe 2018 |
| CES07157 | Otostigmini | *Digitipes jonesii* | JN004039 | JN003928 | JN003983 | Neyyar Wildlife Sanctuary, Thiruvananthapuram District, Kerala, India | Joshi and Karanth 2011 |
| CES07158 | Otostigmini | *Digitipes jonesii* | JX531698 | JX531828 | JX531777 | Neyyar Wildlife Sanctuary, Thiruvananthapuram District, Kerala, India | Joshi and Karanth 2011 |
| CES07159 | Otostigmini | *Rhysida aspinosa* | MK273209 | MK273316 | MK273437 | Neyyar Wildlife Sanctuary, Thiruvananthapuram District, Kerala, India | Joshi and Edgecombe 2018 |
| CES07160 | Otostigmini | *Digitipes jonesii* | JN004040 | JN003929 | JN003984 | Neyyar Wildlife Sanctuary, Thiruvananthapuram District, Kerala, India | Joshi and Karanth 2011 |
| CES07161 | Otostigmini | *Digitipes jonesii* | JN004041 | JN003930 | JN003985 | Neyyar Wildlife Sanctuary, Thiruvananthapuram District, Kerala, India | Joshi and Karanth 2011 |
| CES07162 | Otostigmini | *Digitipes jonesii* | JN004042 | JN003931 | JN003986 | Neyyar Wildlife Sanctuary, Thiruvananthapuram District, Kerala, India | Joshi and Karanth 2011 |
| CES07166 | Otostigmini | *Digitipes jonesii* | JN004043 | JN003932 | JN003987 | Peppara Wildlife Sanctuary, Thiruvananthapuram District, Kerala, India | Joshi and Karanth 2011 |
| CES07168 | Otostigmini | *Digitipes jonesii* | JX531699 | JX531829 | - | Peppara Wildlife Sanctuary, Thiruvananthapuram District, Kerala, India | Joshi and Karanth 2012 |
| CES07171 | Otostigmini | *Digitipes jonesii* | JN004045 | JN003934 | JN003989 | Peppara Wildlife Sanctuary, Thiruvananthapuram District, Kerala, India | Joshi and Karanth 2011 |
| CES07174 | Otostigmini | *Digitipes jonesii* | JN004047 | JN003936 | JN003991 | Peppara Wildlife Sanctuary, Thiruvananthapuram District, Kerala, India | Joshi and Karanth 2011 |
| CES07180 | Otostigmini | *Rhysida pazhuthara* | JN004047 | JN003936 | JN003991 | Ponmudi,Thiruvananthapuram District, Kerala, India | Joshi and Karanth 2011 |
| CES07183 | Otostigmini | *Digitipes jonesii* | JX531700 | JX531830 | JX531778 | Ponmudi,Thiruvananthapuram District, Kerala, India | Joshi and Karanth 2012 |
| CES07196 | Otostigmini | *Digitipes jonesii* | JX531701 | JX531831 | JX531779 | Ponmudi Reserve Forest, Thiruvananthapuram District, Kerala, India | Joshi and Karanth 2012 |
| CES07197 | Otostigmini | *Digitipes jonesii* | JX531702 | JX531832 | JX531780 | Ponmudi Reserve Forest, Thiruvananthapuram District, Kerala, India | Joshi and Karanth 2012 |
| CES07198 | Otostigmini | *Digitipes jonesii* | JX531703 | JX531833 | JX531781 | Ponmudi Reserve Forest, Thiruvananthapuram District, Kerala, India | Joshi and Karanth 2012 |
| CES07210 | Otostigmini | *Digitipes barnabasi* | JX531704 | JX531834 | JX531782 | Bopdev ghat, Pune District, Maharashtra, India | Joshi and Karanth 2012 |
| CES07215 | Otostigmini | *Digitipes barnabasi* | JX531705 | JX531835 | JX531783 | Castle-rock, Uttara Kannada district, Karnatka, India | Joshi and Karanth 2011 |
| CES07219 | Otostigmini | *Digitipes jangii* | JN004048 | JN003937 | JN003992 | Anashi-Dandeli Tiger Reserve, Karwar District, Karnataka, India | Joshi and Karanth 2012 |
| CES07223 | Otostigmini | *Digitipes jangii* | JN004049 | JN003938 | JN003993 | Anashi-Dandeli Tiger Reserve, Karwar District, Karnataka, India | Joshi and Karanth 2012 |
| CES07224 | Otostigmini | *Rhysida lewisi* | JN004023 | JN003912 | JN003968 | Anashi-Dandeli Tiger Reserve, Karwar District, Karnataka, India | Joshi and Karanth 2011 |
| CES07226 | Otostigmini | *Digitipes jangii* | JN004050 | JN003939 | JN003994 | Anashi-Dandeli Tiger Reserve, Karwar District, Karnataka, India | Joshi and Karanth 2012 |
| CES07230 | Otostigmini | *Digitipes jangii* | JN004051 | JN003940 | JN003995 | Anashi-Dandeli Tiger Reserve, Karwar District, Karnataka, India | Joshi and Karanth 2012 |
| CES07233 | Otostigmini | *Digitipes jangii* | JN004052 | JN003941 | JN003996 | Kumta, Uttara Kannada District, Karnataka, India | Joshi and Karanth 2012 |
| CES07237 | Otostigmini | *Digitipes barnabasi* | JX531706 | JX531836 | - | Kumta, Uttara Kannada District, Karnataka, India | Joshi and Karanth 2012 |
| CES07239 | Otostigmini | *Digitipes jangii* | MK273217 | MK273326 | MK273444 | Kumta, Uttara Kannada District, Karnataka, India | Joshi and Edgecombe 2018 |
| CES07244 | Otostigmini | *Digitipes barnabasi* | JX531707 | JX531837 | JX531784 | Dughsagar Water Falls, Goa-Karnataka border, India | Joshi and Karanth 2012 |
| CES07283 | Otostigmini | *Digitipes barnabasi* | MK273230 | MK273340 | MK273456 | Pune, Maharashtra, India | Joshi and Edgecombe 2018 |
| CES07284 | Otostigmini | *Digitipes barnabasi* | JX531708 | JX531838 | JX531785 | Pune, Maharashtra, India | Joshi and Karanth 2012 |
| CES07288 | Otostigmini | *Digitipes jangii* | JX531709 | JX531839 | JX531786 | Talacauvery Reserve Forest, Coorg district, Karnataka, India | Joshi and Karanth 2012 |
| CES08907 | Otostigmini | *Digitipes jangii* | JX531710 | JX531840 | JX531787 | Kurinjal, Kudremukh National Park, India | Joshi and Karanth 2012 |
| CES08911 | Otostigmini | *Digitipes barnabasi* | JX531711 | JX531841 | JX531788 | Kurinjal, Kudremukh National Park, India | Joshi and Karanth 2012 |
| CES08912 | Otostigmini | *Digitipes jangii* | MK273237 | MK273347 | - | Kurinjal, Kudremukh National Park, India | Joshi and Edgecombe 2018 |
| CES08913 | Otostigmini | *Digitipes nudus* | MK273238 | MK273348 | - | Kurinjal, Kudremukh National Park, India | Joshi and Edgecombe 2018 |
| CES08915 | Otostigmini | *Digitipes jangii* | JX531713 | JX531843 | JX531789 | Kurinjal, Kudremukh National Park, India | Joshi and Karanth 2012 |
| CES08922 | Otostigmini | *Digitipes jangii* | JX531715 | JX531845 | JX531791 | Tadoli, Kudremukh National Park, India | Joshi and Karanth 2012 |
| CES08930 | Otostigmini | *Digitipes jangii* | JX531716 | JX531846 | - | Bombay point, Mahabaleshwar Reserve Forest, Maharashtra, India | Joshi and Karanth 2012 |
| CES08932 | Otostigmini | *Digitipes barnabasi* | JX531717 | JX531847 | - | Bombay point, Mahabaleshwar Reserve Forest, Maharashtra, India | Joshi and Karanth 2012 |
| CES08947 | Otostigmini | *Ethmostigmus sahyadrensis* | MH908721 | MH908691 | MH908708 | Amboli, Sindhudurg Dsitrict, Maharashtra, India | Joshi and Edgecombe 2018 |
| CES08951 | Asanadini | *Asanada* sp. | JN004014 | JN003903 | JN003959 | Amboli, Sindhudurg Dsitrict, Maharashtra, India | Joshi and Karanth 2011 |
| CES08953 | Otostigmini | *Digitipes barnabasi* | JX531718 | JX531848 | - | Amboli, Sindhudurg Dsitrict, Maharashtra, India | Joshi and Karanth 2012 |
| CES08957 | Otostigmini | *Digitipes barnabasi* | JX531719 | JX531849 | JX531792 | Amboli, Sindhudurg Dsitrict, Maharashtra, India | Joshi and Karanth 2012 |
| CES08960 | Otostigmini | *Digitipes coonoorensis* | JX531720 | JX531850 | JX531793 | Pambadum Shola National Park, Kerala, India | Joshi and Karanth 2012 |
| CES08961 | Otostigmini | *Digitipes nudus* | MK273245 | MK273357 | MK273467 | Pambadum Shola National Park, Kerala, India | Joshi and Edgecombe 2018 |
| CES08980 | Otostigmini | *Digitipes coonoorensis* | JX531721 | JX531851 | JX531794 | Ervikulam National Park, Kerala, India | Joshi and Karanth 2012 |
| CES08982 | Otostigmini | *Digitipes jonesii* | JX531722 | JX531852 | JX531795 | Ervikulam National Park, Kerala, India | Joshi and Karanth 2012 |
| CES08987 | Otostigmini | *Digitipes coonoorensis* | JX531723 | JX531853 | JX531796 | Ervikulam National Park, Kerala, India | Joshi and Karanth 2012 |
| CES08990 | Otostigmini | *Digitipes jonesii* | JX531724 | JX531854 | JX531797 | Ervikulam National Park, Kerala, India | Joshi and Karanth 2012 |
| CES08992 | Otostigmini | *Digitipes coonoorensis* | JX531725 | JX531855 | JX531798 | Ervikulam National Park, Kerala, India | Joshi and Karanth 2012 |
| CES08994 | Otostigmini | *Digitipes coonoorensis* | JX531726 | JX531856 | - | Ervikulam National Park, Kerala, India | Joshi and Karanth 2012 |
| CES08996 | Otostigmini | *Digitipes jonesii* | JX531727 | JX531857 | JX531799 | Thattekkad Bird Sanctuary, Kerala, India | Joshi and Karanth 2012 |
| CES08997 | Otostigmini | *Digitipes jonesii* | JX531728 | JX531858 | JX531800 | Thattekkad Bird Sanctuary, Kerala, India | Joshi and Karanth 2012 |
| CES091005 | Otostigmini | *Digitipes jonesii* | JX531730 | JX531860 | - | Peechi-Vazhani Wildlife Sanctuary, Kerala, India | Joshi and Karanth 2012 |
| CES091006 | Otostigmini | *Digitipes jonesii* | JX531731 | JX531861 | JX531802 | Peechi-Vazhani Wildlife Sanctuary, Kerala, India | Joshi and Karanth 2012 |
| CES091008 | Otostigmini | *Digitipes jonesii* | JX531732 | JX531862 | JX531803 | Peechi-Vazhani Wildlife Sanctuary, Kerala, India | Joshi and Karanth 2011 |
| CES091011 | Otostigmini | *Ethmostigmus praveeni* | MH908722 | MH908686 | MH908703 | Kudremukh National Park, India | Joshi and Edgecombe 2018 |
| CES091016 | Otostigmini | *Digitipes barnabasi* | JX531734 | JX531864 | JX531805 | Talacauvery, Kodagu district, Karnataka, India | Joshi and Karanth 2012 |
| CES091017 | Otostigmini | *Digitipes barnabasi* | JX531735 | JX531865 | JX531806 | Talacauvery, Kodagu district, Karnataka, India | Joshi and Karanth 2012 |
| CES091020 | Otostigmini | *Digitipes jangii* | JX531736 | JX531866 | JX531807 | Bisale Ghat Reserve Forest, Karnataka, India | Joshi and Karanth 2012 |
| CES091029 | Otostigmini | *R.* *konda* | MK273251 | MK273367 | MK273475 | Sitakund, Mayurbhanj district, Odisha, India | Joshi and Edgecombe 2018 |
| CES091033 | Otostigmini | *Digitipes barnabasi* | JX531738 | JX531868 | JX531808 | Periyar Tiger Reserve, Kerala, India | Joshi and Karanth 2012 |
| CES091037 | Otostigmini | *Digitipes nudus* | JX531739 | JX531869 | JX531809 | Vellimalai, Periyar Tiger Reserve, Kerala, India | Joshi and Karanth 2012 |
| CES091038 | Otostigmini | *Digitipes nudus* | JX531740 | JX531870 | - | Vellimalai, Periyar Tiger Reserve, Kerala, India | Joshi and Karanth 2012 |
| CES091039 | Otostigmini | *Digitipes barnabasi* | JX531741 | JX531871 | JX531810 | Vellimalai, Periyar Tiger Reserve, Kerala, India | Joshi and Karanth 2012 |
| CES091040 | Otostigmini | *Rhysida trispinosa* | MK273255 | MK273371 | MK273479 | Nayanoor, Andhra Pradesh, India | Joshi and Edgecombe 2018 |
| CES091047 | Otostigmini | *Digitipes jonesii* | JX531742 | JX531872 | JX531811 | Parambikulam Tiger Reserve, Kerala, India | Joshi and Karanth 2012 |
| CES091049 | Otostigmini | *Digitipes jonesii* | JX531744 | JX531874 | JX531812 | Parambikulam Tiger Reserve, Kerala, India | Joshi and Karanth 2012 |
| CES091057 | Otostigmini | *Digitipes jonesii* | JX531745 | JX531875 | JX531813 | Sholayar Reserve Forest, Kerala, India | Joshi and Karanth 2012 |
| CES091062 | Otostigmini | *Digitipes jonesii* | JX531746 | JX531876 | JX531814 | Vazhachal Reserve Forest, Kerala, India | Joshi and Karanth 2012 |
| CES091065 | Otostigmini | *Ethmostigmus agasthyamalaiensis* | MH908726 | MH908695 | MH908710 | Parambikulam Tiger Reserve, Kerala, India | Joshi and Edgecombe 2018 |
| CES091073 | Otostigmini | *Digitipes barnabasi* | JX531747 | JX531877 | - | Parakkadavu, Shendurney Wildlife Sanctuary, Kerala, India | Joshi and Karanth 2012 |
| CES091086 | Otostigmini | *Digitipes jonesii* | JX531748 | JX531878 | JX531815 | Pandimatte, Shendurney Wildlife Sanctuary, Kollam district, Kerala, India | Joshi and Karanth 2012 |
| CES091087 | Otostigmini | *Digitipes jonesii* | JX531749 | JX531879 | JX531816 | Pandimatte, Shendurney Wildlife Sanctuary, Kollam district, Kerala, India | Joshi and Karanth 2012 |
| CES091088 | Otostigmini | *Digitipes coonoorensis* | JX531750 | JX531880 | JX531817 | Pandimatte, Shendurney Wildlife Sanctuary, Kollam district, Kerala, India | Joshi and Karanth 2012 |
| CES091089 | Otostigmini | *Digitipes jonesii* | JX531751 | JX531881 | - | Pandimatte, Shendurney Wildlife Sanctuary, Kollam district, Kerala, India | Joshi and Karanth 2012 |
| CES091090 | Otostigmini | *Digitipes jonesii* | JX531752 | JX531882 | - | Pandimatte, Shendurney Wildlife Sanctuary, Kollam district, Kerala, India | Joshi and Karanth 2012 |
| CES091091 | Otostigmini | *Digitipes jonesii* | JX531753 | JX531883 | - | Pandimatte, Shendurney Wildlife Sanctuary, Kollam district, Kerala, India | Joshi and Karanth 2012 |
| CES091096 | Otostigmini | *Digitipes jonesii* | JX531754 | JX531884 | - | Pandimatte, Shendurney Wildlife Sanctuary, Kollam district, Kerala, India | Joshi and Karanth 2012 |
| CES091304 | Otostigmini | *Digitipes jonesii* | JX531755 | JX531885 | - | Achankovil Reserve Forest, Pathanamthitta district, Kerala, India | Joshi and Karanth 2012 |
| CES091305 | Otostigmini | *Digitipes jonesii* | JX531756 | JX531886 | - | Achankovil Reserve Forest, Pathanamthitta district, Kerala, India | Joshi and Karanth 2012 |
| CES091310 | Otostigmini | *Digitipes jonesii* | JX531757 | JX531887 | - | Vellithode, Periyar Tiger Reserve, Kerala, India | Joshi and Karanth 2012 |
| CES091313 | Otostigmini | *Digitipes jonesii* | MK273272 | MK273394 | MK273494 | Vellithode, Periyar Tiger Reserve, Kerala, India | Joshi and Edgecombe 2018 |
| CES091315 | Otostigmini | *Digitipes jonesii* | MK273274 | JX531888 | - | Vellithode, Periyar Tiger Reserve, Kerala, India | Joshi and Karanth 2012 |
| CES091318 | Otostigmini | *Digitipes jonesii* | JX531760 | JX531890 | - | Vellithode, Periyar Tiger Reserve, Kerala, India | Joshi and Karanth 2012 |
| CES091319 | Otostigmini | *Digitipes jonesii* | JX531761 | JX531891 | - | Vellithode, Periyar Tiger Reserve, Kerala, India | Joshi and Karanth 2012 |
| CES091322 | Otostigmini | *Digitipes jonesii* | JX531762 | JX531892 | - | Periyar Tiger Reserve, Kerala, India | Joshi and Karanth 2012 |
| CES091324 | Otostigmini | *Digitipes jonesii* | JX531763 | JX531893 | - | Kottavasal Reserve Forest, Kerala, India | Joshi and Karanth 2012 |
| CES091325 | Otostigmini | *Digitipes jonesii* | JX531764 | JX531894 | JX531818 | Kottavasal Reserve Forest, Kerala, India | Joshi and Karanth 2012 |
| CES091326 | Otostigmini | *Digitipes jonesii* | JX531765 | JX531895 | - | Kottavasal Reserve Forest, Kerala, India | Joshi and Karanth 2012 |
| CES091334 | Otostigmini | *Digitipes coonoorensis* | JX531769 | JX531899 | JX531821 | Siruvani Reserve Forest, Kerala, India | Joshi and Karanth 2012 |
| CES091335 | Otostigmini | *Ethmostigmus agasthyamalaiensis* | MH908729 | MH908697 | MH908712 | Kottavasal Reserve Forest, Kerala, India | Joshi and Edgecombe 2018 |
| CES091341 | Otostigmini | *Digitipes barnabasi* | JX531770 | JX531900 | - | Pasikada, Aachankovil Reserve Forest, Kerala, India | Joshi and Karanth 2012 |
| CES091345 | Otostigmini | *Digitipes barnabasi* | JX531772 | JX531902 | JX531823 | Bhimashankar, Maharashtra, India | Joshi and Karanth 2012 |
| CES091348 | Otostigmini | *Digitipes barnabasi* | JX531773 | JX531903 | JX531824 | Bhimashankar, Maharashtra, India | Joshi and Karanth 2012 |
| CES091353 | Otostigmini | *Digitipes barnabasi* | JX531774 | JX531904 | JX531825 | Bhimashankar, Maharashtra, India | Joshi and Karanth 2012 |
| CES091355 | Otostigmini | *Digitipes barnabasi* | JX531775 | JX531905 | JX531826 | Bhimashankar, Maharashtra, India | Joshi and Karanth 2012 |
| CES091357 | Otostigmini | *Digitipes barnabasi* | MK273278 | MK273401 | - | Bhimashankar, Maharashtra, India | Joshi and Edgecombe 2018 |
| CES091431 | Otostigmini | *Digitipes barnabasi* | - | MN535701 | - | Sakaleshapur, Karnataka, India | Joshi et al. 2020 |
| CES091450 | Otostigmini | *Digitipes coonoorensis* | MK273298 | MK273422 | MK273516 | Mesapulimalai, Kerala, India | Joshi and Edgecombe 2018 |
| CES091461 | Otostigmini | *Ethmostigmus* n. sp. | - | MK296124 | MK296126 | Bellinchep, Tripura, India | Joshi and Edgecombe 2018b |
| CES091469 | Otostigmini | *Digitipes barnabasi* | - | MN535704 | - | Coorg, Karnataka, India | Joshi et al. 2020 |
| CES181505 | Otostigmini | *Digitipes jonesii* | MN535713 | MN535705 | MN535721 | Ponmudi Reserve Forest, Thiruvananthapuram District, Kerala, India | Joshi et al. 2020 |
| CES181513 | Otostigmini | *Digitipes jonesii* | MN535714 | MN535706 | MN535722 | IISER campus, Thiruvananthapuram District, Kerala, India | Joshi et al. 2020 |
| CES181516 | Otostigmini | *Digitipes jonesii* | MN535715 | MN535707 | MN535723 | Thenmala, Kollam district, Kerala, India | Joshi et al. 2020 |
| CES181530 | Otostigmini | *Digitipes coonoorensis* | MN535717 | MN535709 | MN535725 | Waynad, Kerala, India | Joshi et al. 2020 |
| CES181538 | Otostigmini | *Digitipes jonesii* | MN535718 | MN535710 | MN535726 | Waynad, Kerala, India | Joshi et al. 2020 |
| MCZ DNA100454 | Otostigmini | *Alipes crotalus* | AY288742 | AY288720 | HM453273 | Swaziland | Vahtera *et al*. 2012 |
| MCZ DNA106771 | Otostigmini | *Alipes grandidieri* | KF676514 | KF676473 | KF676371 | Tanzania | Vahtera *et al*. 2013 |
| MCZ DNA104719 | Asanadini | *Asanada socotrana* | HQ402541 | HQ402490 | HQ402524 | Senegal | Vahtera *et al*. 2012 |
| MCZ DNA103951 | Scolopendrini | *Cormocephalus aurantiipes* | HQ402543 | HQ402492 | KF676388 | Western Australia, Australia | Vahtera *et al*. 2012 |
| MCZ DNA106751 | Scolopendrini | *Cormocephalus nitidus* | KF676533 | KF676492 | KF676397 | KwaZulu-Natal, South Africa | Vahtera *et al*. 2013 |
| CUMZ<THA>:00233 | Otostigmini | *Digitipes kalewaensis* | KP204114 | KP204110 | - | Kale District, Sagaing Division, Myanmar | Siriwut *et al*. 2015a |
| CUMZ<THA>:00234 | Otostigmini | *Digitipes kalewaensis* | KP204115 | KP204111 | - | Kale District, Sagaing Division, Myanmar | Siriwut *et al*. 2015a |
| CUMZ<THA>:00235 | Otostigmini | *Digitipes kalewaensis* | KP204116 | KP204112 | - | Kale District, Sagaing Division, Myanmar | Siriwut *et al*. 2015a |
| CUMZ<THA>:00236 | Otostigmini | *Digitipes kalewaensis* | KP204117 | KP204113 | - | Kale District, Sagaing Division, Myanmar | Siriwut *et al*. 2015a |
| CUMZ 00517.1 | Otostigmini | *Digitipes* sp. | MF167811 | MF167744 | MF167878 | Phra Cave, Ban Namyen, Myanmar | Siriwut *et al*. 2018 |
| CUMZ 00517.2 | Otostigmini | *Digitipes* sp. | MF167812 | MF167745 | MF167879 | Phra Cave, Ban Namyen, Myanmar | Siriwut *et al*. 2018 |
| WAM T115537 | Otostigmini | *Ethmostigmus curtipes* | KF676515 | KF676474 | KF676372 | Western Australia, Australia | Vahtera *et al*. 2013 |
| MCZ DNA106500 | Otostigmini | *Otostigmus angusticeps* | KF676509 | - | KF676365 | Finisterre Mountains, Papua New Guinea | Vahtera *et al*. 2013 |
| CUMZ 00522 | Otostigmini | *Otostigmus multidens* | MF167778 | MF167711 | MF167845 | Kao-Nui Park, Ruttaphume, Songkhla, Thailand | Siriwut *et al*. 2018 |
| MCZ DNA104870 | Otostigmini | *Otostigmus rugulosus* | HQ402550 | HQ402499 | HQ402534 | Wangthong, Phitsanulok, Thailand | Vahtera *et al*. 2012 |
| MCZ DNA106491 | Otostigmini | *Otostigmus scaber* | KF676513 | KF676471 | KF676369 | Ilan County, Taiwan | Vahtera *et al*. 2013 |
| CMUZ 00232 | Otostigmini | *Otostigmus spinosus* | MF167783 | MF167716 | MF167850 | Ban na ka som, Attapue, Laos | Siriwut *et al*. 2018 |
| CUMZ 00532 | Otostigmini | *Otostigmus sulcipes* | MF167803 | MF167736 | MF167870 | Huai Hong Khrai, Chiang Mai, Thailand | Siriwut *et al*. 2018 |
| MCZ DNA | Scolopendrini | *Scolopendra cingulata* | HM453310 | HM453220 | AF000782 | Catalunya, Spain | Vahtera *et al*. 2013 |
| CCMB400 | Otostigmini | *Digitipes barnabasi* | + | + | + | Talacauvery WLS, Karnataka, India | This study |
| CCMB404 | Otostigmini | *Digitipes barnabasi* | + | + | + | Talacauvery WLS, Karnataka, India | This study |
| CCMB409 | Otostigmini | *Digitipes barnabasi* | + | + | + | Talacauvery WLS, Karnataka, India | This study |
| CCMB412 | Otostigmini | *Digitipes jangii* | + | + | + | Talacauvery WLS, Karnataka, India | This study |
| CCMB428 | Otostigmini | *Digitipes jangii* | + | + | + | Talacauvery WLS, Karnataka, India | This study |
| CCMB431 | Otostigmini | *Digitipes barnabasi* | + | + | + | Talacauvery WLS, Karnataka, India | This study |
| CCMB433 | Otostigmini | *Digitipes barnabasi* | + | + | + | Talacauvery WLS, Karnataka, India | This study |
| CCMB461 | Otostigmini | *Digitipes barnabasi* | + | + | + | Talacauvery WLS, Karnataka, India | This study |
| CCMB463 | Otostigmini | *Digitipes barnabasi* | + | + | + | Talacauvery WLS, Karnataka, India | This study |
| CCMB475 | Otostigmini | *Digitipes jangii* | + | + | + | Pushpagiri WLS, Karnataka, India | This study |
| CCMB482 | Otostigmini | *Digitipes jangii* | + | + | + | Pushpagiri WLS, Karnataka, India | This study |
| CCMB506 | Otostigmini | *Digitipes barnabasi* | + | + | + | Pushpagiri WLS/Subramanya, Karnataka, India | This study |
| CCMB530 | Otostigmini | *Digitipes barnabasi* | + | + | + | Pushpagiri WLS/Galibeedu, Karnataka, India | This study |
| CCMB603 | Otostigmini | *Digitipes jangii* | + | + | + | Makutta Reserve Forest, Karnataka, India | This study |
| CCMB606 | Otostigmini | *Digitipes jangii* | + | + | + | Makutta Reserve Forest, Karnataka, India | This study |
| CCMB653 | Otostigmini | *Digitipes barnabasi* | + | + | + | Makutta Reserve Forest, Kokka, Karnataka, India | This study |
| CCMB655 | Otostigmini | *Digitipes jangii* | + | + | + | Makutta Reserve Forest, Kokka, Karnataka, India | This study |
| CCMB1326 | Otostigmini | *Digitipes pruthii* | + | + | + | Horsley Hills, Andhra Pradesh, India | This study |
| CCMB1327 | Otostigmini | *Digitipes pruthii* | + | + | + | Horsley Hills, Andhra Pradesh, India | This study |
| CCMB1328 | Otostigmini | *Digitipes pruthii* | + | + | + | Horsley Hills, Andhra Pradesh, India | This study |
| CCMB1329 | Otostigmini | *Digitipes pruthii* | + | + | + | Horsley Hills, Andhra Pradesh, India | This study |
| CCMB1337 | Otostigmini | *Digitipes coonoorensis* | + | + | + | Silent Valley National Park, Kerala, India | This study |
| CCMB1373 | Otostigmini | *Digitipes coonoorensis* | + | + | + | Silent Valley National Park, Kerala, India | This study |
| CCMB1384 | Otostigmini | *Digitipes coonoorensis* | + | + | + | Silent Valley National Park, Kerala, India | This study |
| CCMB1386 | Otostigmini | *Digitipes coonoorensis* | + | + | + | Silent Valley National Park, Kerala, India | This study |
| CCMB1389 | Otostigmini | *Digitipes coonoorensis* | + | + | + | Silent Valley National Park, Kerala, India | This study |
| CCMB1924 | Otostigmini | *Digitipes coonoorensis* | + | + | + | Anamudi Shola National Park, Kerala, India | This study |
| CCMB1926 | Otostigmini | *Digitipes coonoorensis* | + | + | + | Anamudi Shola National Park, Kerala, India | This study |
| CCMB1952 | Otostigmini | *Digitipes coonoorensis* | + | + | + | Anamudi Shola National Park, Kerala, India | This study |
| CCMB1953 | Otostigmini | *Digitipes coonoorensis* | + | + | + | Anamudi Shola National Park, Kerala, India | This study |
| CCMB2148 | Otostigmini | *Digitipes jonesii* | + | + | + | Peechi Wildlife Sanctuary, Kerala, India | This study |

#### S3. Primers details and cycling conditions

| Marker | Amplicon length (bp) | Primer | Reference | Sequence (5’- 3’) | Annealing Temperature (℃) |
| --- | --- | --- | --- | --- | --- |
| COI | 656 | LCO1490 (Forward) | Folmer et al. (1994) | GGTCAACAAATCATAAAGATATTGG | 46 |
|  |  | HCO2198 (Reverse) |  | TAAACTTCAGGGTGACCAAAAAATCA |  |
| 16S rRNA | 531 | 16Sar (Forward) | Xiong and Kocher (1991) | CGCCTGTTTATCAAAAACAT | 49 |
|  |  | 16Sb (Reverse) |  | CTCCGGTTTGAACTCAGATCA |  |
| 28S rRNA | 486 | 28Sa (Forward) | Whiting et al. (1997) | GACCCGTCTTGAAACACGGA | 52 |
| 28S rRNA | 486 | 28Sb (Reverse) |  | TCGGAAGGAACCAGCTAC |  |

#### S4. *Digitipes* Species Delimitation (mPTP)

mPTP analysis using the results of *Digitipes* from Otostigminae phylogeny and COI sequences of *Digitipes* (n=140).

Minimum branch length was set to 0.0008577573, Null model log-likelihood: 627.294627, Best score for multi-coalescent rate: 710.702218

Number of delimited species was 24

| Species | Voucher | Taxa |
| --- | --- | --- |
| 1 | CES07125 | *Digitipes coonoorensis* |
|  | CES07134 | *Digitipes coonoorensis* |
|  | CES07132 | *Digitipes coonoorensis* |
|  | CES07137 | *Digitipes coonoorensis* |
|  | CES07126 | *Digitipes coonoorensis* |
|  | CES091334 | *Digitipes coonoorensis* |
|  | CCMB1384 | *Digitipes coonoorensis* |
|  | CCMB1386 | *Digitipes coonoorensis* |
|  | CCMB1389 | *Digitipes coonoorensis* |
|  | CCMB1373 | *Digitipes coonoorensis* |
| 2 | CES181530 | *Digitipes coonoorensis* |
| 3 | CES07157 | *Digitipes jonesii* |
|  | CES07162 | *Digitipes jonesii* |
|  | CES07174 | *Digitipes jonesii* |
|  | CES07171 | *Digitipes jonesii* |
|  | CES07166 | *Digitipes jonesii* |
|  | CES07197 | *Digitipes jonesii* |
|  | CES07198 | *Digitipes jonesii* |
|  | CES091310 | *Digitipes jonesii* |
|  | CES181505 | *Digitipes jonesii* |
|  | CES181513 | *Digitipes jonesii* |
|  | CES181516 | *Digitipes jonesii* |
|  | CES091086 | *Digitipes jonesii* |
|  | CES091087 | *Digitipes jonesii* |
|  | CES091089 | *Digitipes jonesii* |
|  | CES091091 | *Digitipes jonesii* |
|  | CES091096 | *Digitipes jonesii* |
|  | CES091090 | *Digitipes jonesii* |
|  | CES091304 | *Digitipes jonesii* |
|  | CES091305 | *Digitipes jonesii* |
|  | CES091324 | *Digitipes jonesii* |
|  | CES091325 | *Digitipes jonesii* |
|  | CES091326 | *Digitipes jonesii* |
|  | CES07158 | *Digitipes jonesii* |
|  | CES07160 | *Digitipes jonesii* |
|  | CES07161 | *Digitipes jonesii* |
|  | CES07196 | *Digitipes jonesii* |
| 4 | CES07168 | *Digitipes jonesii* |
|  | CES08996 | *Digitipes jonesii* |
|  | CES08997 | *Digitipes jonesii* |
|  | CES091005 | *Digitipes jonesii* |
|  | CES091006 | *Digitipes jonesii* |
|  | CES091008 | *Digitipes jonesii* |
|  | CES091047 | *Digitipes jonesii* |
|  | CES091049 | *Digitipes jonesii* |
|  | CCMB2148 | *Digitipes jonesii* |
|  | CES091057 | *Digitipes jonesii* |
|  | CES091062 | *Digitipes jonesii* |
|  | CES091313 | *Digitipes jonesii* |
|  | CES091315 | *Digitipes jonesii* |
|  | CES091318 | *Digitipes jonesii* |
|  | CES091319 | *Digitipes jonesii* |
|  | CES091322 | *Digitipes jonesii* |
|  | CES07183 | *Digitipes jonesii* |
| 5 | CCMB1326 | *Digitipes pruthii* |
|  | CCMB1327 | *Digitipes pruthii* |
|  | CCMB1329 | *Digitipes pruthii* |
|  | CCMB1328 | *Digitipes pruthii* |
| 6 | CES08960 | *Digitipes coonoorensis* |
|  | CCMB1924 | *Digitipes coonoorensis* |
|  | CCMB1926 | *Digitipes coonoorensis* |
|  | CCMB1953 | *Digitipes coonoorensis* |
|  | CES091450 | *Digitipes coonoorensis* |
|  | CCMB1952 | *Digitipes coonoorensis* |
|  | CES08994 | *Digitipes coonoorensis* |
|  | CES08980 | *Digitipes coonoorensis* |
|  | CES08987 | *Digitipes coonoorensis* |
|  | CES08992 | *Digitipes coonoorensis* |
| 7 | CES091088 | *Digitipes coonoorensis* |
| 8 | CES08982 | *Digitipes jonesii* |
|  | CES08990 | *Digitipes jonesii* |
| 9 | CES181538 | *Digitipes jonesii* |
|  | CCMB1337 | *Digitipes coonoorensis* |
| 10 | CES07127 | *Digitipes nudus* |
|  | CES07133 | *Digitipes nudus* |
|  | CES08913 | *Digitipes nudus* |
|  | CES08961 | *Digitipes nudus* |
|  | CES091037 | *Digitipes nudus* |
|  | CES091038 | *Digitipes nudus* |
| 11 | CES07219 | *Digitipes jangii* |
|  | CES07239 | *Digitipes jangii* |
|  | CES07233 | *Digitipes jangii* |
| 12 | CES07223 | *Digitipes jangii* |
|  | CES07230 | *Digitipes jangii* |
|  | CES07226 | *Digitipes jangii* |
| 13 | CES07288 | *Digitipes jangii* |
|  | CCMB412 | *Digitipes jangii* |
|  | CCMB428 | *Digitipes jangii* |
|  | CCMB606 | *Digitipes jangii* |
|  | CCMB655 | *Digitipes jangii* |
|  | CCMB603 | *Digitipes jangii* |
| 14 | CES08912 | *Digitipes jangii* |
| 15 | CES08907 | *Digitipes jangii* |
|  | CES08915 | *Digitipes jangii* |
|  | CES091020 | *Digitipes jangii* |
|  | CCMB475 | *Digitipes jangii* |
|  | CCMB482 | *Digitipes jangii* |
|  | CES08922 | *Digitipes jangii* |
|  | CES08930 | *Digitipes jangii* |
| 16 | CES07130 | *Digitipes barnabasi* |
|  | CES07138 | *Digitipes barnabasi* |
|  | CES08911 | *Digitipes barnabasi* |
|  | CCMB404 | *Digitipes barnabasi* |
|  | CCMB433 | *Digitipes barnabasi* |
|  | CCMB530 | *Digitipes barnabasi* |
|  | CES091017 | *Digitipes barnabasi* |
|  | CCMB400 | *Digitipes barnabasi* |
|  | CES091469 | *Digitipes barnabasi* |
|  | CES091073 | *Digitipes barnabasi* |
|  | CES091341 | *Digitipes barnabasi* |
| 17 | CES07210 | *Digitipes barnabasi* |
|  | CES07283 | *Digitipes barnabasi* |
|  | CES07284 | *Digitipes barnabasi* |
|  | CES091348 | *Digitipes barnabasi* |
|  | CES091353 | *Digitipes barnabasi* |
|  | CES091357 | *Digitipes barnabasi* |
|  | CES091355 | *Digitipes barnabasi* |
|  | CES091345 | *Digitipes barnabasi* |
| 18 | CES07215 | *Digitipes barnabasi* |
|  | CES07244 | *Digitipes barnabasi* |
|  | CES07237 | *Digitipes barnabasi* |
| 19 | CES08932 | *Digitipes barnabasi* |
| 20 | CES08953 | *Digitipes barnabasi* |
|  | CES08957 | *Digitipes barnabasi* |
| 21 | CES091016 | *Digitipes barnabasi* |
|  | CCMB461 | *Digitipes barnabasi* |
|  | CCMB431 | *Digitipes barnabasi* |
|  | CCMB506 | *Digitipes barnabasi* |
|  | CES091431 | *Digitipes barnabasi* |
|  | CCMB409 | *Digitipes barnabasi* |
|  | CCMB653 | *Digitipes barnabasi* |
|  | CCMB463 | *Digitipes barnabasi* |
| 22 | CES091033 | *Digitipes barnabasi* |
|  | CES091039 | *Digitipes barnabasi* |
| 23 | CUMZ<THA>:00233 | *Digitipes kalewaensis* |
|  | CUMZ<THA>:00235 | *Digitipes kalewaensis* |
|  | CUMZ<THA>:00236 | *Digitipes kalewaensis* |
|  | CUMZ<THA>:00234 | *Digitipes kalewaensis* |
| 24 | CUMZ 00517.1 | *Digitipes sp.* |
|  | CUMZ 00517.2 | *Digitipes sp.* |

#### 5. ASAP analysis using the COI sequences of *Digitipes* (n=117).

Species delimitation using ASAP on a subset of the COI dataset (117 sequences, 530 bp) recovering 35 putative species in the best-ranked partition (ASAP score = 6.5, distance threshold = 0.0556).

We report both cluster size and uniformity to indicate the clarity and reliability of ASAP‑assigned clusters.

ASAP cluster composition: “Uniform” = all individuals in the cluster belong to the same species; “Mostly uniform” = the cluster is dominated by one species, but may include one or a few individuals of a different species (outliers marked as grey); “Mixed” = the cluster contains multiple species with no clear dominant species.

Cluster size class: Clusters with ≤ 3 individuals are classified as “Small”; all others as “Well represented”

| ASAP cluster | Voucher | Taxa | Cluster composition | Cluster Size |
| --- | --- | --- | --- | --- |
| 1 | CES07125 | *Digitipes coonoorensis* | Uniform | Large |
|  | CES07134 | *Digitipes coonoorensis* |  |  |
|  | CES07132 | *Digitipes coonoorensis* |  |  |
|  | CES07126 | *Digitipes coonoorensis* |  |  |
|  | CES07137 | *Digitipes coonoorensis* |  |  |
|  | CCMB1384 | *Digitipes coonoorensis* |  |  |
|  | CCMB1386 | *Digitipes coonoorensis* |  |  |
|  | CCMB1389 | *Digitipes coonoorensis* |  |  |
|  | CCMB1373 | *Digitipes coonoorensis* |  |  |
| 2 | CES181530 | *Digitipes coonoorensis* | Uniform | Small |
| 3 | CES07157 | *Digitipes jonesii* | Mostly Uniform | Large |
|  | CES07162 | *Digitipes jonesii* |  |  |
|  | CES07158 | *Digitipes jonesii* |  |  |
|  | CES181513 | *Digitipes jonesii* |  |  |
|  | CES07160 | *Digitipes jonesii* |  |  |
|  | CES181516 | *Digitipes jonesii* |  |  |
|  | CES07161 | *Digitipes jonesii* |  |  |
|  | CES181505 | *Digitipes jonesii* |  |  |
|  | CES07198 | *Digitipes jonesii* |  |  |
|  | CES07196 | *Digitipes jonesii* |  |  |
|  | CES07171 | *Digitipes jonesii* |  |  |
|  | CES07166 | *Digitipes jonesii* |  |  |
|  | CES07197 | *Digitipes jonesii* |  |  |
|  | CES091324 | *Digitipes jonesii* |  |  |
|  | CES091326 | *Digitipes jonesii* |  |  |
|  | CES091310 | *Digitipes jonesii* |  |  |
|  | CES091341 | *Digitipes barnabasi* |  |  |
|  | CES091304 | *Digitipes jonesii* |  |  |
|  | CES091305 | *Digitipes jonesii* |  |  |
|  | CES07174 | *Digitipes jonesii* |  |  |
|  | CES091086 | *Digitipes jonesii* |  |  |
|  | CES091089 | *Digitipes jonesii* |  |  |
|  | CES091091 | *Digitipes jonesii* |  |  |
|  | CES091096 | *Digitipes jonesii* |  |  |
|  | CES091090 | *Digitipes jonesii* |  |  |
| 4 | CES07168 | *Digitipes jonesii* | Uniform | Small |
|  | CES07183 | *Digitipes jonesii* |  |  |
| 5 | CES08996 | *Digitipes jonesii* | Uniform | Small |
|  | CES08997 | *Digitipes jonesii* |  |  |
| 6 | CES091005 | *Digitipes jonesii* | Uniform | Small |
|  | CES091006 | *Digitipes jonesii* |  |  |
| 7 | CES091008 | *Digitipes jonesii* | Mostly Uniform | Large |
|  | CCMB475 | *Digitipes jangii* |  |  |
|  | CCMB482 | *Digitipes jangii* |  |  |
|  | CES08907 | *Digitipes jangii* |  |  |
|  | CES08915 | *Digitipes jangii* |  |  |
|  | CES08922 | *Digitipes jangii* |  |  |
| 8 | CES091047 | *Digitipes jonesii* | Mostly Uniform | Small |
|  | CES091049 | *Digitipes jonesii* |  |  |
|  | CCMB2148 | *Digitipes jonesii* |  |  |
| 9 | CES091062 | *Digitipes jonesii* | Uniform | Small |
| 10 | CES091322 | *Digitipes jonesii* | Uniform | Small |
| 11 | CCMB1326 | *Digitipes pruthii* | Uniform | Large |
|  | CCMB1327 | *Digitipes pruthii* |  |  |
|  | CCMB1329 | *Digitipes pruthii* |  |  |
|  | CCMB1328 | *Digitipes pruthii* |  |  |
| 12 | CES08960 | *Digitipes coonoorensis* | Uniform | Large |
|  | CCMB1952 | *Digitipes coonoorensis* |  |  |
|  | CES08994 | *Digitipes coonoorensis* |  |  |
|  | CES091450 | *Digitipes coonoorensis* |  |  |
|  | CES08980 | *Digitipes coonoorensis* |  |  |
|  | CES091088 | *Digitipes coonoorensis* |  |  |
|  | CES08987 | *Digitipes coonoorensis* |  |  |
|  | CES08992 | *Digitipes coonoorensis* |  |  |
| 13 | CES08982 | *Digitipes jonesii* | Uniform | Small |
|  | CES08990 | *Digitipes jonesii* |  |  |
| 14 | CES181538 | *Digitipes jonesii* | Mixed | Small |
|  | CCMB1337 | *Digitipes coonoorensis* |  |  |
| 15 | CES07127 | *Digitipes nudus* | Uniform | Small |
|  | CES07133 | *Digitipes nudus* |  |  |
| 16 | CES08913 | *Digitipes nudus* | Uniform | Small |
| 17 | CES08961 | *Digitipes nudus* |  | Small |
| 18 | CES091037 | *Digitipes nudus* |  | Small |
| 19 | CES07219 | *Digitipes jangii* | Uniform | Small |
|  | CES07239 | *Digitipes jangii* |  |  |
|  | CES07233 | *Digitipes jangii* |  |  |
| 20 | CES07223 | *Digitipes jangii* | Uniform | Small |
|  | CES07230 | *Digitipes jangii* |  |  |
|  | CES07226 | *Digitipes jangii* |  |  |
| 21 | CES07288 | *Digitipes jangii* | Uniform | Large |
|  | CCMB412 | *Digitipes jangii* |  |  |
|  | CCMB428 | *Digitipes jangii* |  |  |
|  | CCMB603 | *Digitipes jangii* |  |  |
|  | CCMB655 | *Digitipes jangii* |  |  |
|  | CCMB606 | *Digitipes jangii* |  |  |
| 22 | CES08912 | *Digitipes jangii* | Uniform | Small |
| 23 | CES091020 | *Digitipes jangii* | Uniform | Small |
| 24 | CES08930 | *Digitipes jangii* | Uniform | Small |
| 25 | CES07138 | *Digitipes barnabasi* | Uniform | Small |
| 26 | CCMB404 | *Digitipes barnabasi* | Uniform | Small |
|  | CCMB433 | *Digitipes barnabasi* |  |  |
|  | CCMB530 | *Digitipes barnabasi* |  |  |
| 27 | CCMB400 | *Digitipes barnabasi* | Uniform | Small |
| 28 | CES091073 | *Digitipes barnabasi* | Uniform | Small |
| 29 | CES07210 | *Digitipes barnabasi* | Uniform | Large |
|  | CES07283 | *Digitipes barnabasi* |  |  |
|  | CES07284 | *Digitipes barnabasi* |  |  |
|  | CES091348 | *Digitipes barnabasi* |  |  |
|  | CES091357 | *Digitipes barnabasi* |  |  |
|  | CES091355 | *Digitipes barnabasi* |  |  |
| 30 | CES07215 | *Digitipes barnabasi* | Uniform | Small |
|  | CES07244 | *Digitipes barnabasi* |  |  |
|  | CES07237 | *Digitipes barnabasi* |  |  |
| 31 | CES08932 | *Digitipes barnabasi* | Uniform | Small |
| 32 | CES08953 | *Digitipes barnabasi* | Uniform | Small |
|  | CES08957 | *Digitipes barnabasi* |  |  |
| 33 | CCMB431 | *Digitipes barnabasi* | Uniform | Large |
|  | CCMB506 | *Digitipes barnabasi* |  |  |
|  | CCMB409 | *Digitipes barnabasi* |  |  |
|  | CCMB463 | *Digitipes barnabasi* |  |  |
|  | CCMB653 | *Digitipes barnabasi* |  |  |
| 34 | CUMZ<THA>:00233 | *Digitipes kalewaensis* | Uniform | Large |
|  | CUMZ<THA>:00236 | *Digitipes kalewaensis* |  |  |
|  | CUMZ<THA>:00234 | *Digitipes kalewaensis* |  |  |
|  | CUMZ<THA>:00235 | *Digitipes kalewaensis* |  |  |
| 35 | CUMZ 00517.1 | *Digitipes sp.* | Uniform | Small |
|  | CUMZ 00517.2 | *Digitipes sp.* |  |  |

S6. 10 best ASAP partitions for the *Digitipes* COI dataset (sequence length = 530 bp), ranked by ASAP score

35 putative species in the best-ranked partition (ASAP score = 6.5). The lower the ASAP score, the better the model.

| Rank | Distance | No. of species | ASAP score |
| --- | --- | --- | --- |
| 1 | 0.0556 | 35 | 6.5 (Best Ranked Partition) |
| 2 | 0.0690 | 30 | 7.5 |
| 3 | 0.0647 | 31 | 8.5 |
| 4 | 0.0076 | 74 | 9 |
| 5 | 0.0865 | 25 | 10 |
| 6 | 0.0927 | 23 | 12 |
| 7 | 0.0095 | 70 | 12 |
| 8 | 0.0171 | 55 | 13 |
| 9 | 0.0506 | 38 | 14 |
| 10 | 0.0797 | 26 | 14.5 |

S7. Fossil Calibration for Divergence Time Estimation with starBEAST

| **Calibration type** | **References** | **Position** | **Clade** | **Age Interval (mya)** | **Fossil** |
| --- | --- | --- | --- | --- | --- |
| fossil | Wolfe et al. 2016; Fernández et al. 2016 | Crown | Pleurostigmophora | 382.6 - 382.8 | *Devonobius delta* |
| fossil | Wolfe et al. 2016; Fernández et al. 2016 | Crown | Epimorpha | 306.9 - 307.1 | *Mazoscolopendra richardsoni* |
| fossil | Wolfe et al. 2016; Joshi and Edgecombe 2019 | Crown | Scolopendridae | 112.6 - 307 | *Cratoraricrus oberlii* |
| secondary | Joshi et al. 2020 | Crown | Otostignminae | 189.9 - 282.5 | - |
| secondary | Joshi et al. 2020 | Crown | *Rhysida immarginata* complex | 62.5 - 150.9 | - |
| secondary | Joshi et al. 2020 | Crown | *Rhysida longipes* complex | 48.3- 148.9 | - |
| secondary | Joshi and Edgecombe 2019 | Crown | Ethmostigmus | 58.6 - 129.7 | - |

#### S8. Species-delimitation (mPTP) support and geography to determine emerging clades in *Digitipes*

mPTP species‑delimitation support is the proportion of MCMC samples in which a node is treated as a split between two species, rather than as variation within a single species. It ranges from 0 to 1.

Area codes:

S (Southern Western Ghats), C (Central Western Ghats), N (Northern Western Ghats), E (Eastern Ghats), A (Mainland South-east Asia)

mPTP delimitation support ranges from 0 - 1

| mPTP species comparison | Taxa comparison and geography | mPTP  Delimitation support | Emerging  Clades |
| --- | --- | --- | --- |
| 1 - 2 | *D. coonoorensis (C) - D. coonoorensis (C)* | 1 | *D. coonoorensis (C)* clade 1 |
| 3 - 4 | *D. jonesii (S) - D. jonesii (S)* | 1 | *D. jonesii (S)* |
| (3 & 4) - 5 | *D. jonesii (S) & D. chhotanii (S) - D. pruthii (E)* | 1 | *D. pruthii (E)* |
| (3 to 5) - (6 & 7) | *D. jonesii (S) & D. chhotanii (S) & D. pruthii (E) - D. coonoorensis (S)* | 1 | *-* |
| 6 - 7 | *D. coonoorensis (S) - D. coonoorensis (S)* | 0.99 | *D. coonoorensis (S)* clade 2 |
| (3 to 7) - (8 & 9) | *D. jonesii (S) & D. chhotanii (S) & D. pruthii (E) & D. coonoorensis (S) - D. jonesii (S) & D. jonesii(C) & D. coonoorensis (C)* | 1 | *-* |
| 8 - 9 | *D. jonesii (S) - D. coonoorensis (C) & D. jonesii (C)* | 0.97 | *D. sp. A (S & C)*; comprising of  *D. jonesii (S) & D. coonoorensis (C) & D. jonesii (C)* |
| (3 to 9) - (1 & 2) | *D. jonesii (S) & D. chhotanii (S) & D. pruthii (E) & D. coonoorensis (S) & D. sp. A (S & C) - D. coonoorensis (C)* | 1 | - |
| (1 to 9) - 10 | *D. coonoorensis (C) & D. jonesii (S) & D. chhotanii (S) & D. pruthii (E) & D. coonoorensis (S) & D. sp. A (S & C) - D. nudus (S) & D. nudus (C)* | 1 | *D. nudus (S & C)* |
| (1 to 10) - (11 to 15) | *D. coonoorensis (C) & D. jonesii (S) & D. chhotanii (S) & D. pruthii (E) & D. coonoorensis (S) & D. sp. A (S & C) & D. nudus (S & C) - D. jangii (C) & D. jangii (N)* | 1 | - |
| (11 & 12) - (13 & 14) | *D. jangii (C) - D. jangii (C)* | 1 | - |
| 11 - 12 | *D. jangii (C) - D. jangii (C)* | 0.94 | *D. jangii* *(C)* clade 3 |
| 13 - 14 | *D. jangii (C) - D. jangii (C)* | 0.95 | *D. jangii* *(C)* clade 1 |
| (11 to 14) - 15 | *D. jangii (C) - D. jangii (C) & D. jangii (N)* | 1 | *D. jangii (C & N)* clade 2 |
| (1 to 15) - (16 to 24) | *D. coonoorensis (C) & D. jonesii (S) & D. chhotanii (S) & D. pruthii (E) & D. coonoorensis (S) & D. sp. A (S & C) & D. nudus (S & C) & D. jangii (C & N) - D. barnabasi (C) & D. barnabasi (S) & D. barnabasi (N) & D. kalewaensis (A) & D. sp.(A)* | 1 | - |
| 16 - (17 to 21) | *D. barnabasi (C) & D. barnabasi (S) - D. barnabasi (N) & D. barnabasi (C)* | 1 | *D. barnabasi (C & S)* clade 2 |
| (17 & 18) - 19 | *D. barnabasi (N) - D. barnabasi (N)* | 1 | *D. barnabasi (N)* clade 4 |
| 20 - 21 | *D. barnabasi (N) - D. barnabasi (C)* | 1 | *D. barnabasi (N & C)* clade 1 |
| (16 to 21) - 22 | *D. barnabasi (C) & D. barnabasi (S) & D. barnabasi (N) - D. barnabasi (S)* | 1 | *D. barnabasi (C & S)* clade 3 |
| (17 to 19) - (20 & 21) | *D. barnabasi (N) - D. barnabasi (N) & D. barnabasi (C)* | 1 | - |
| 17 - 18 | *D. barnabasi (N) - D. barnabasi (N)* | 0.99 | - |
| (16 to 22) - (23 & 24) | *D. barnabasi (C) & D. barnabasi (S) & D. barnabasi (N) - D. kalewaensis (A) & D. sp.(A)* | 1 | - |
| 23 - 24 | *D. kalewaensis (A) & D. sp.(A)* | 1 | *D. kalewaensis (A), D. sp.(A)* |

S9. BioGeoBEARS analysis - Area Codes, Time Periods and Dispersal Multiplier Assignment

Area Codes: S (Southern Western Ghats), C (Central Western Ghats), N (Northern Western Ghats), E (Eastern Ghats), A (Mainland South-east Asia)

| **Time Points** | **Event** |
| --- | --- |
| 150 | Time prior to estimated crown age of *Digitipes* |
| 65 | Deccan volcanism |
| 55 | First ever transient land connection with Asia |
| 20 | Beginning of climatic shifts that caused aridification |
| Present |  |

| **Dispersal Multiplier** | **Condition** |
| --- | --- |
| 1 | Contiguous areas |
| 0.75 | Adjacent areas with moderate connectivity but has a barrier |
| 0.5 | Moderately separated areas with intermediate connections, or separated by another area/two or more landmasses |
| 0.0001 | Well-separated areas by water/large terrestrial barrier |

S10. BioGeoBEARS analysis - Dispersal matrices

Dispersal matrices at each time strata for A2 analysis given below.

For the A1 analysis (null hypothesis), all values were set to 1. (Supplementary material 3, point 2.)

| **150 (mya)** | **No connection with South-east Asia** | | | | |
| --- | --- | --- | --- | --- | --- |
| Areas | S | C | N | E | A |
| S | 1 | 0.75 | 0.5 | 1 | 0.00001 |
| C | 0.75 | 1 | 1 | 1 | 0.00001 |
| N | 0.5 | 1 | 1 | 1 | 0.00001 |
| E | 1 | 1 | 1 | 1 | 0.00001 |
| A | 0.00001 | 0.00001 | 0.00001 | 0.00001 | 1 |
| **65 (mya)** | **Deccan Volcanism, no connection with Southeast Asia** | | | | |
| Areas | S | C | N | E | A |
| S | 1 | 0.75 | 0.5 | 1 | 0.00001 |
| C | 0.75 | 1 | 1 | 1 | 0.00001 |
| N | 0.5 | 1 | 1 | 1 | 0.00001 |
| E | 1 | 1 | 1 | 1 | 0.00001 |
| A | 0.00001 | 0.00001 | 0.00001 | 0.00001 | 1 |
| **55 (mya)** | **First connection with Southeast Asia; India acts as a biotic ferry** | | | | |
| Areas | S | C | N | E | A |
| S | 1 | 0.75 | 0.5 | 1 | 0.5 |
| C | 0.75 | 1 | 1 | 1 | 0.5 |
| N | 0.5 | 1 | 1 | 1 | 0.5 |
| E | 1 | 1 | 1 | 1 | 0.5 |
| A | 0.5 | 0.5 | 0.5 | 0.5 | 1 |
| **20 (mya)** | **Global heating, Aridification in the subcontinent, Out-of India dispersals reduced** | | | | |
| Areas | S | C | N | E | A |
| S | 1 | 0.75 | 0.5 | 0.5 | 0.00001 |
| C | 0.75 | 1 | 0.75 | 0.5 | 0.00001 |
| N | 0.5 | 0.75 | 1 | 0.5 | 0.00001 |
| E | 0.5 | 0.5 | 0.5 | 1 | 0.00001 |
| A | 0.00001 | 0.00001 | 0.00001 | 0.00001 | 1 |
| **Present** | **Dispersal barrier (Indo-Gangetic plains) between India and Southeast Asia** | | | | |
| Areas | S | C | N | E | A |
| S | 1 | 0.75 | 0.5 | 0.5 | 0.00001 |
| C | 0.75 | 1 | 0.75 | 0.5 | 0.00001 |
| N | 0.5 | 0.75 | 1 | 0.5 | 0.00001 |
| E | 0.5 | 0.5 | 0.5 | 1 | 0.00001 |
| A | 0.00001 | 0.00001 | 0.00001 | 0.00001 | 1 |
